# Integrated multi-omics analysis reveals divergent molecular responses in Palmer amaranth (*Amaranthus palmeri*) biotypes susceptible and resistant to glyphosate

**DOI:** 10.1101/2025.07.20.665775

**Authors:** Pawanjit Kaur Sandhu, Rohit Kumar, Vijay Nandula, Nishanth Tharayil

## Abstract

Environmental stress triggers coordinated changes across genetic, transcriptomic, proteomic, and metabolomic levels in plants, yet the extent of synchronization across these *omic* layers remains underexplored. We captured transcriptomic, proteomic and metabolomic perturbation of glyphosate-resistant (GR) and glyphosate-susceptible (GS) Palmer amaranth (*Amaranthus palmeri*) biotypes 24 hours after herbicide treatment, quantifying 30,371 transcripts, 5,606 proteins, and 220 metabolites. Glyphosate perturbed threefold more transcripts and proteins in GS than in GR and caused the accumulation of shikimate intermediates in both biotypes. In GS, glyphosate severely disrupted primary metabolism, including photosynthesis and carbon fixation, leading to a collapse of energy production and impairment of phenylpropanoid and terpenoid biosynthesis, compromising defense and detoxification. In contrast, GR maintained cellular homeostasis, with minimal perturbation in carbon metabolism and upregulation of detoxifying pathways, indicating metabolic rerouting. Integrated multi-omics analysis captured stress responses hidden from single-omic analysis, including elevated glutathione metabolism, perturbation of the phenylpropanoid pathway and elevated raffinose family oligosaccharide metabolism in GR, and perturbation of taurine-hypotaurine metabolism in GS. Transcript and protein changes were broadly correlated, but GS exhibited signs of translational inhibition under glyphosate stress, indicating reduced protein synthesis. These findings reveal pervasive perturbation of glyphosate beyond the shikimate pathway within 24 hours after exposure, and underscore the importance of multi-omics integration to elucidate complex stress responses in plants.

## INTRODUCTION

Environmental stressors induce a plethora of changes in plants at genetic, transcriptomic, proteomic, and metabolomic levels (Patti et al., 2012; Haak et al., 2017; Gimenez et al., 2018; Sun and Zhou, 2018). Recent advancements in next-generation sequencing and mass spectrometry have allowed us to study these modifications by acquiring high-throughput ‘omics’ data for genes (genomics), transcripts (transcriptomics), proteins (proteomics), and metabolites (metabolomics) (Misra et al., 2019; Jamil et al., 2020). These approaches are often used in isolation to decipher the cellular physiology of plant stress responses. The integration of global omics approaches could be a more effective strategy for investigating stress-induced responses in plants that are coordinated across multiple levels of cellular regulation (Haak et al., 2017; Gimenez et al., 2018; Sun and Zhou, 2018). So far, the integrated omics studies focusing on stress responses have been limited to model plants and crops (Soda et al., 2015; Meena et al., 2017; Schwartz, 2020). However, some of the most robust stress adaptive strategies are exhibited by ruderal plants, including extremophiles, agricultural weeds, and invasive species (Clements and Ditommaso, 2011; Oh et al., 2012; Délye et al., 2013; Gaines et al., 2020). Thus, the extension of multi-omics studies, including the integration of transcriptomics, proteomics, and metabolomics approaches, to non-model species could provide a better understanding of the stress-induced perturbations and mitigation mechanisms in plants.

Herbicide resistance in agricultural weeds is a complex phenomenon regulated by the multiplicity and diversity of resistance mechanisms (Délye et al., 2013; Heap and Duke, 2018; Heap, 2023). To date, resistance to two-thirds of the known herbicidal modes of action has been reported across 273 weed species worldwide (Heap, 2025). Similar to other stress adaptations, herbicide resistance mechanisms in weeds exhibit pleiotropic effects (Dyer, 2018), which range from modifications in target enzyme specificity (Vila-Aiub et al., 2019) to the reduction in the overall fitness (Vila-Aiub et al., 2009). Although studying individual components of the central dogma (genes, transcripts, proteins, metabolites, etc.) has provided critical insights into the mechanisms per se (Délye et al., 2013; Gaines et al., 2020), an integrated knowledge of how the resistance is regulated at the molecular level is currently lacking.

Palmer amaranth (*Amaranthus palmeri* [S. Wats]) is a noxious weed that causes yield losses ranging from 50-90% in row crop production systems (corn, soybean, and cotton) of the southeastern United States (Ward et al., 2013). Mainstay for the control of Palmer amaranth has been the herbicide glyphosate, which blocks the shikimate pathway in plants, leading to the depletion of aromatic amino acids and, thus, a slower plant death. Resistance to glyphosate in Palmer amaranth was reported in 2006 in Georgia (Culpepper et al., 2006) and since then has spread to forty-eight states across the US (Heap, 2025). The primary resistance mechanism to glyphosate in Plamner amaranth is the amplification of the gene that codes for the target enzyme, 5-Enolpyruvylshikimate-3-phosphate synthase (EPSPS) (Gaines et al., 2010). Genomics and molecular biology studies have shown that the glyphosate-resistant (GR) Palmer amaranth possesses a constitutively higher abundance of EPSPS transcript, protein, and enzyme activity (Gaines et al., 2010; 2011). This suggests that the physiology of GR Palmer amaranth would remain unaffected after glyphosate exposure as excess EPSPS would ensure a non-disruptive functioning of shikimate pathway (Gaines et al., 2010; 2011). On the other hand, the whole plant physiology and metabolomics studies have consistently shown glyphosate-induced disruption of cellular physiology across GR biotypes, (Nandula et al., 2012; Maroli et al., 2015; Fernández-Escalada et al., 2016; 2017; 2019; Sandhu et al., 2023), including photosynthesis (Campbell et al. 1976; Vaughn and Duke 1986) and respiration (Kielak et al., 2011), that are less directly connected to disruption of the shikimate pathway. These physiological perturbations point to the prevalence of off-target toxicity of glyphosate and/or a potential time lag in the functionalization of the resistance mechanism after the herbicide exposure in this species. The simultaneous profiling of perturbations at the transcriptome, proteome and metabolome levels could provide insights into glyphosate-induced changes at the molecular level.

This study aimed to investigate the multi-level differential response induced by glyphosate in two biotypes of Palmer amaranth that differ in their susceptibility to the herbicide glyphosate. The primary objectives were to discern the unique and shared responses to glyphosate exposure in these biotypes at various levels of cellular physiology, including the global transcriptome, proteome, and metabolome. Additionally, the study aimed to elucidate how these molecular changes interact and contribute to the overall phenotype of glyphosate resistance or susceptibility. This integrative approach helps in understanding the broader physiological and biochemical impacts of glyphosate on Palmer amaranth and could inform effective management strategies for this troublesome weed.

## RESULTS

### Global changes in transcriptome, proteome, and metabolome induced after glyphosate treatment

The alignment of ∼44 million (98.59%; Q20 > 98.57%; Q30 > 95.54%; 42.44% GC Content; Table S1) reads to Palmer amaranth genome resulted in 19463 and 19852 expressed transcripts for GR and GS, respectively. Across the biotypes, the control and glyphosate-treated shared 90% of the transcripts [Figure 1A]. Across the biotypes and treatments, the proteomic analysis resulted in 604198 peptide spectrum matches (PSM), 37923 peptide groups, and 5050 master proteins [Figure 1B]. Metabolomic analysis identified 220 primary and secondary metabolites in Palmer amaranth. Eighty metabolites were identified from gas chromatography-quadrupole-time of flight (GC-QToF) analysis, based on retention index (RI) and fragmentation pattern matches with standards or Kovats library (confidence level 1 and 2, respectively; Schmansky et al. 2014) (Figure 1C, Table S2), while 140 metabolites were annotated at confidence levels 2 and 3 from ultrahigh pressure liquid chromatography-tandem mass-spectrometry (UHPLC-MS/MS) analysis [Figure 1D, Table S3]. The identified metabolites included primary metabolites (amino acids, sugars, organic acids, fatty acids) and secondary metabolites (phenylpropanoids, terpenoids, fatty acyl glycosides, etc.) [Figure 1C, 1D; Table S2, S3]. Since a significant proportion of mass features (>70%) from UHPLC-MS/MS analysis were unidentified and were annotated into ClassyFire superclass and class (Djoumbou Feunang et al., 2016) using CANOPUS (Dührkop et al., 2021) [Table S4]. The Partial Lease Square – Discriminatory Analysis (PLS-DA) showed a clear distinction between the biotypes and treatments in all the omics datasets, indicating that glyphosate produced treatment effects in both GS and GR at all measured levels of cellular physiology, including transcriptome, proteome, primary, and secondary metabolome [Figure S1].

**Figure 1.**
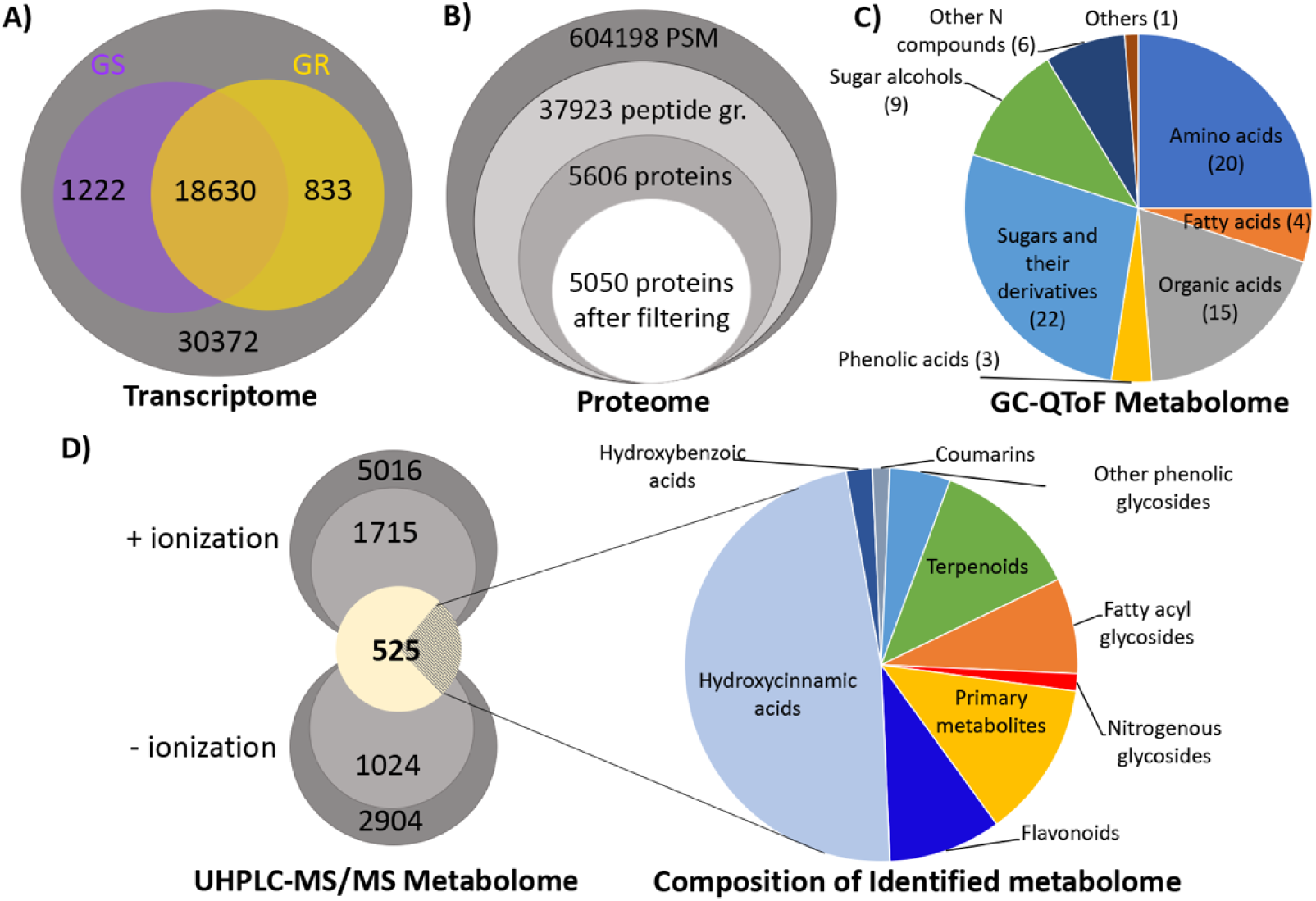
General overview of global Transcriptome (A.), Proteome (B.), Metabolome from GC-QToF (C.), and UHPLC-MS/MS (D.) of Palmer Amaranth captured in the present study.

### Glyphosate-induced changes in transcriptome and proteome of GS and GR biotypes

Glyphosate-induced changes in transcript expression were translated for 99% of the measured proteins [Table S5, S6, S7, S8]. Overall transcript abundance showed a positive association (r = 0.53-0.59) with protein abundance across the biotypes and treatments [Figure 2A; Figure S2; Table S8]. Compared to their respective control, when treated with glyphosate, thrice the number of transcripts and proteins were differentially expressed (DE) in GS (3349 DE transcripts and 226 DE proteins) than in GR (1028 DE transcripts and 71 DE proteins) [Figure 2B; Table S5, S6, S7]. Glyphosate treatment resulted in a similar number of transcripts being upregulated and downregulated in GS (1588 up vs. 1761 down), whereas the transcriptome of glyphosate-treated GR was mostly upregulated [Figure 2B]. Across the biotypes, glyphosate stress-induced changes in proteome were driven by protein upregulation, with three times and five times the number of proteins upregulated compared to the down-regulated proteins in GR and GS, respectively [Figure 2B; Table S5, S6, S7]. Further, more than half of DE transcripts and proteins in GR were also disrupted in GS (806 transcripts and 41 proteins; Figure 2B), capturing a potential similarity in the stress response of these two biotypes. A fraction of the DE transcripts showed a change in the corresponding protein (Figure 2B), although the change in protein abundance was lower than the change in transcript abundance [Table S5, S6, S7]. Thirty of these transcript-protein pairs were common across the biotypes. Interestingly, the downregulated profile was mostly biotype-specific at both transcript and protein levels, with only one common DE protein and 207 DE transcripts across the two biotypes [Figure 2B].

**Figure 2.**
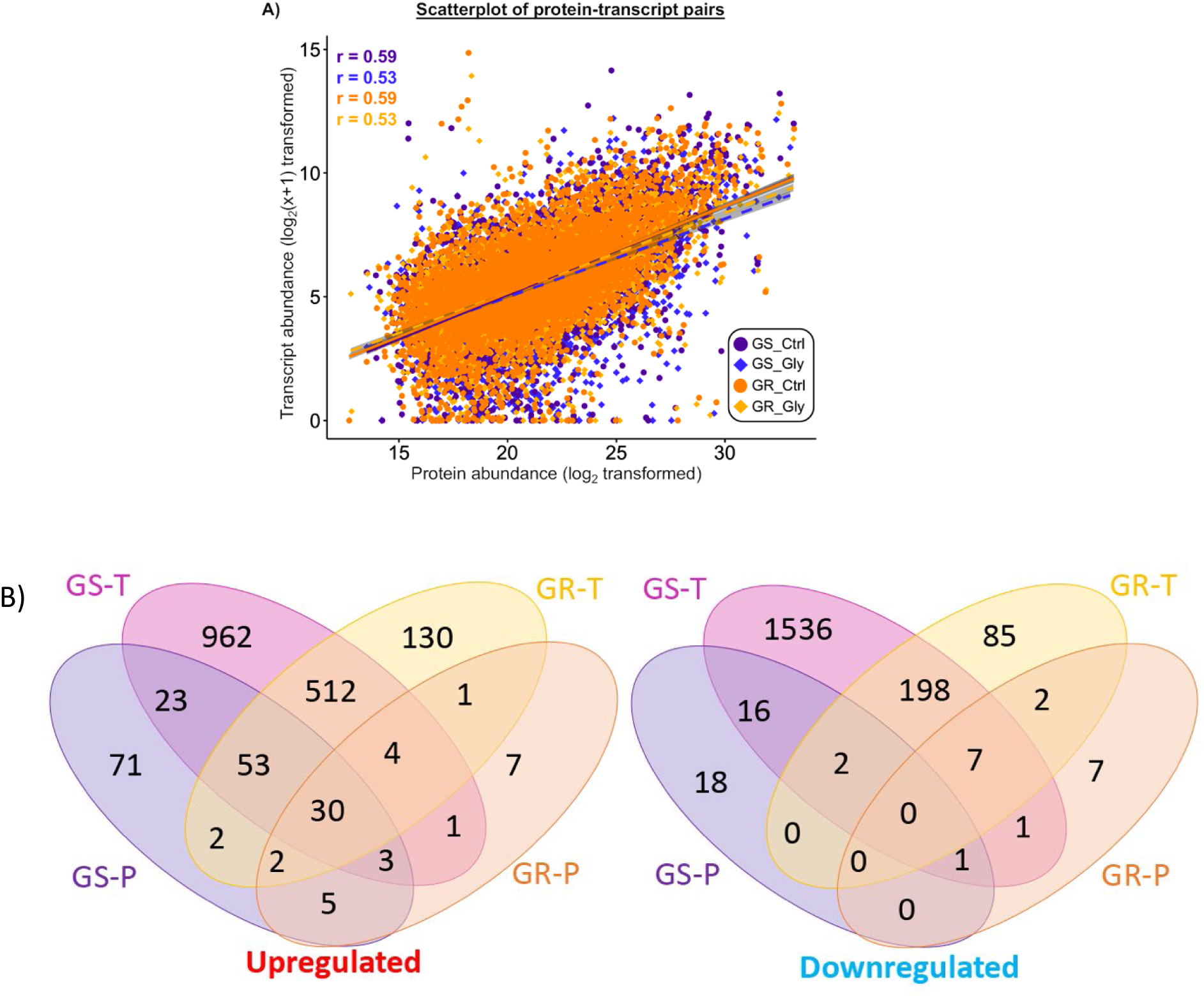
Scatterplot of protein and transcript abundance (A) and venn diagram showing the upregulated and downregulated transcripts (T) and proteins (P) in Glyphosate-susceptible (GS-) and resistant (GR-) biotype (glyphosate-treated vs control) of Amaranthus palmeri (B). Each dot represents mean value. GS-T: Differentially expressed (DE) transcripts in GS-bitoype; GS-P: DE proteins in GS-biotype; GR-T: DE transcripts in GR-biotype; GR-P: DE proteins in GR-biotype.

### Functional enrichment of differentially expressed transcripts and proteins

For functional evaluation, Gene Ontology (GO) enrichment analysis was carried out for DE transcripts and proteins [Figure 3, 4, 5; Table S9]. Compared to the respective controls, GO enrichment showed notable glyphosate-induced upregulation of abscisic acid (ABA)-activated signaling pathway, signaling receptor activity, and protein phosphatase inhibitor activity at both the transcript and protein levels across the biotypes (Figure 3, 5). GS and GR also shared upregulation of an additional twenty-three GO terms in the transcriptome related to protein phosphorylation, regulation of transcription (DNA-templated), transmembrane transport, oxidoreductase activity, nutrient reservoir activity, and transferase activity (MF), among others [Figure 3]. The transcriptome of glyphosate-treated GS showed a specific upregulation of translation and ribosome biogenesis, while GR had a specific upregulation of calcium and ammonium transmembrane transport, carbohydrate metabolic process, defense response to fungi and bacteria, chitin catabolic process, and flavin adenine dinucleotide binding [Figure 4].

**Figure 3.**
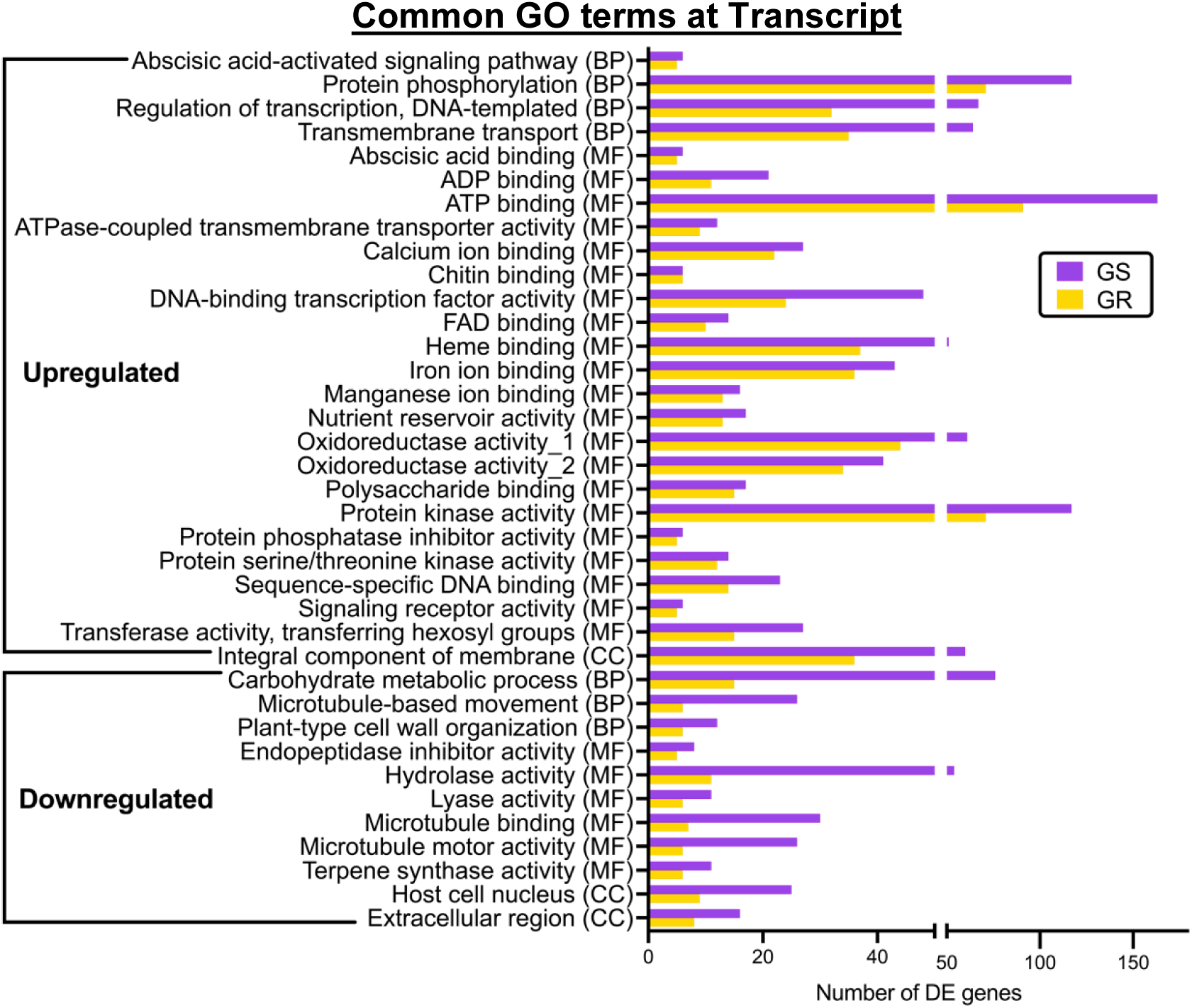
Commonly enriched GO terms (p<0.05; FDR < 0.05) at the transcript level between the glyphosate-treated GS-and GR-biotype of A. palmeri. The Y-axis represents the number of differentially expressed (DE) genes in each category.

**Figure 4.**
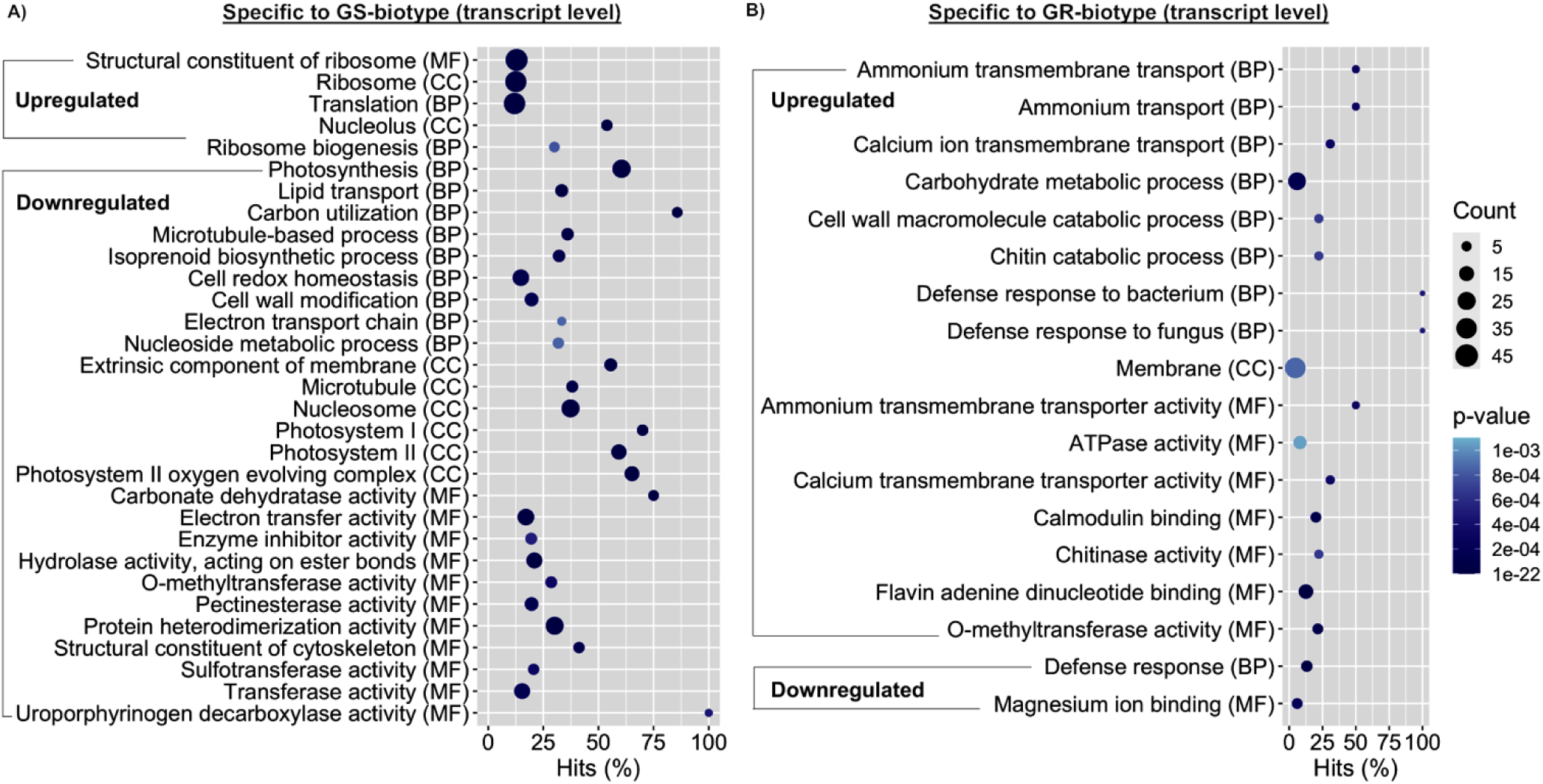
GO-terms significantly enriched (p<0.05; FDR <0.05) at transcript level that are specific to the glyphosate-treated A). GS-and B) GR-biotype. The size of the circle represents the actual number of genes that are differentially expressed (DE) in each GO category while the Y-axis represents the percentage of the genes (% Hits) that are DE in a GO category. p-value is represented by the shades of blue. BP: Biological process; CC: Cellular component; MF: Molecular function.

**Figure 5.**
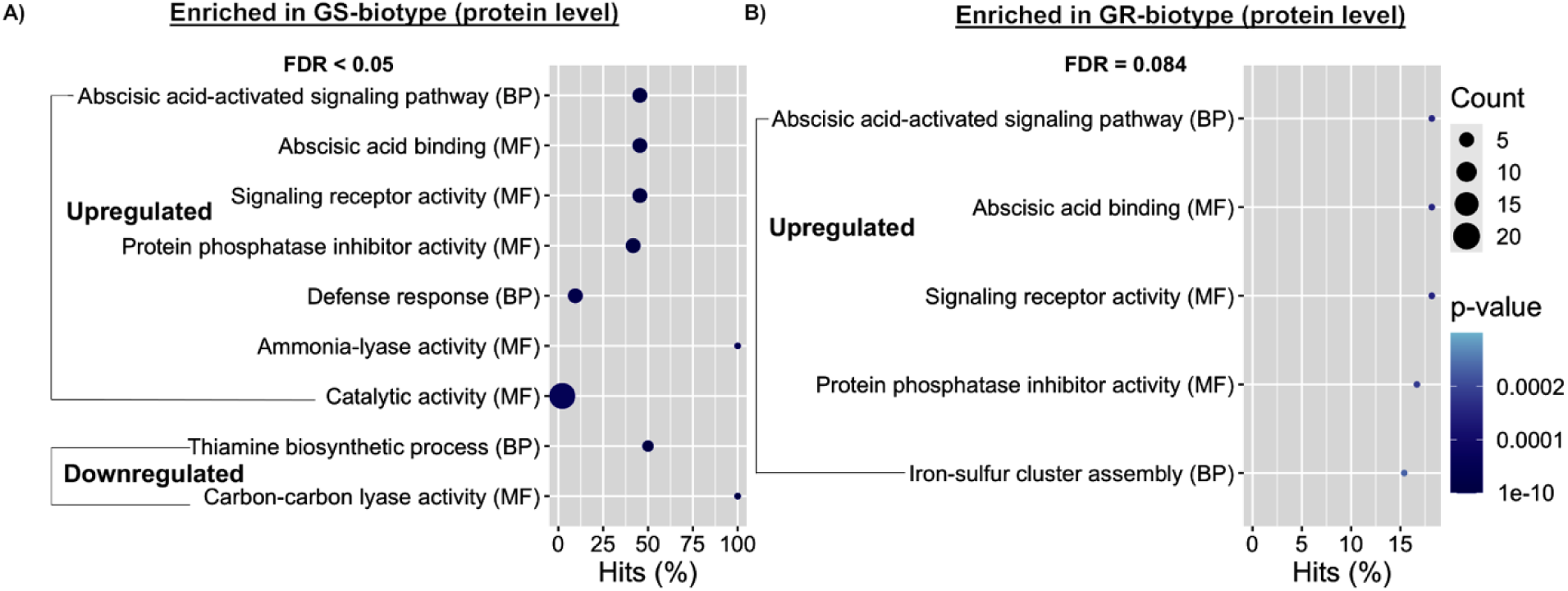
GO-terms significantly enriched (p <0.05) at protein level in the glyphosate-treated A). GS-and B) GR-biotype. The size of the circle represents the actual number of genes that are differentially expressed (DE) in each GO category while the Y-axis represents the percentage of the genes (% Hits) that are DE in a GO category. p-value is represented by the shades of blue. BP: Biological process; CC: Cellular component; MF: Molecular function.

Interestingly, GS also observed a glyphosate-induced downregulation of photosynthesis, lipid transport, carbon utilization, isoprenoid biosynthesis, redox homeostasis, cell wall modification, electron transport chain, and nucleoside metabolic process at the transcript level and thiamine biosynthetic process and carbon-carbon lyase activity at the protein level that was not observed in the GR compared to the control [Figure 3, 4, 5].

### Glyphosate induced changes in the metabolome of GS and GR biotypes

Glyphosate caused significant disruption of primary metabolism in GS, consistent with the changes in expression of transcript and protein [Figure 6A; Table S10]. We observed a reduction in the abundance of three tricarboxylic acid (TCA) cycle intermediates while eleven non-aromatic amino acids (including their derivatives), glucose, and intermediates of the shikimate pathway (shikimic acid, 3-dehydroshikimic acid, and shikimate-3-phosphate) were accumulated in the glyphosate-treated GS [Figure 6A]. Contrastingly, in the GR, the effect on primary metabolism was limited to the accumulation of intermediates of the shikimate pathway, and two amino acid metabolites and the reduction of one TCA cycle intermediate [Figure 6; Table S10]. Further, the glyphosate-induced increase of shikimate pathway intermediates was higher in GS compared to GR [Table S10]. In contrast to transcript, protein and primary metabolites, the effect of glyphosate on secondary metabolites was more significant in GR than GS [Figure 6B; Table S11] and the differences were pronounced in the downregulated profile. Compared to respective controls, glyphosate-treated GR showed a glyphosate-induced reduction in 79 secondary metabolites compared to 21 in the GS. The downregulated secondary metabolite profile, specific to the GR, was dominated by quercetin glycosides, ferulic acid derivatives [Figure 6A] and triterpenoid glycosides [Figure S3]. On the other hand, a similar number of secondary metabolites accumulated across the biotypes (41 upregulated metabolites in GR vs 47 in GS) [Figure 6B; Table S11], including hydroxybenzoic acid derivatives [Figure 6A], organic acid derivatives and oxygen-containing compounds.

**Figure 6.**
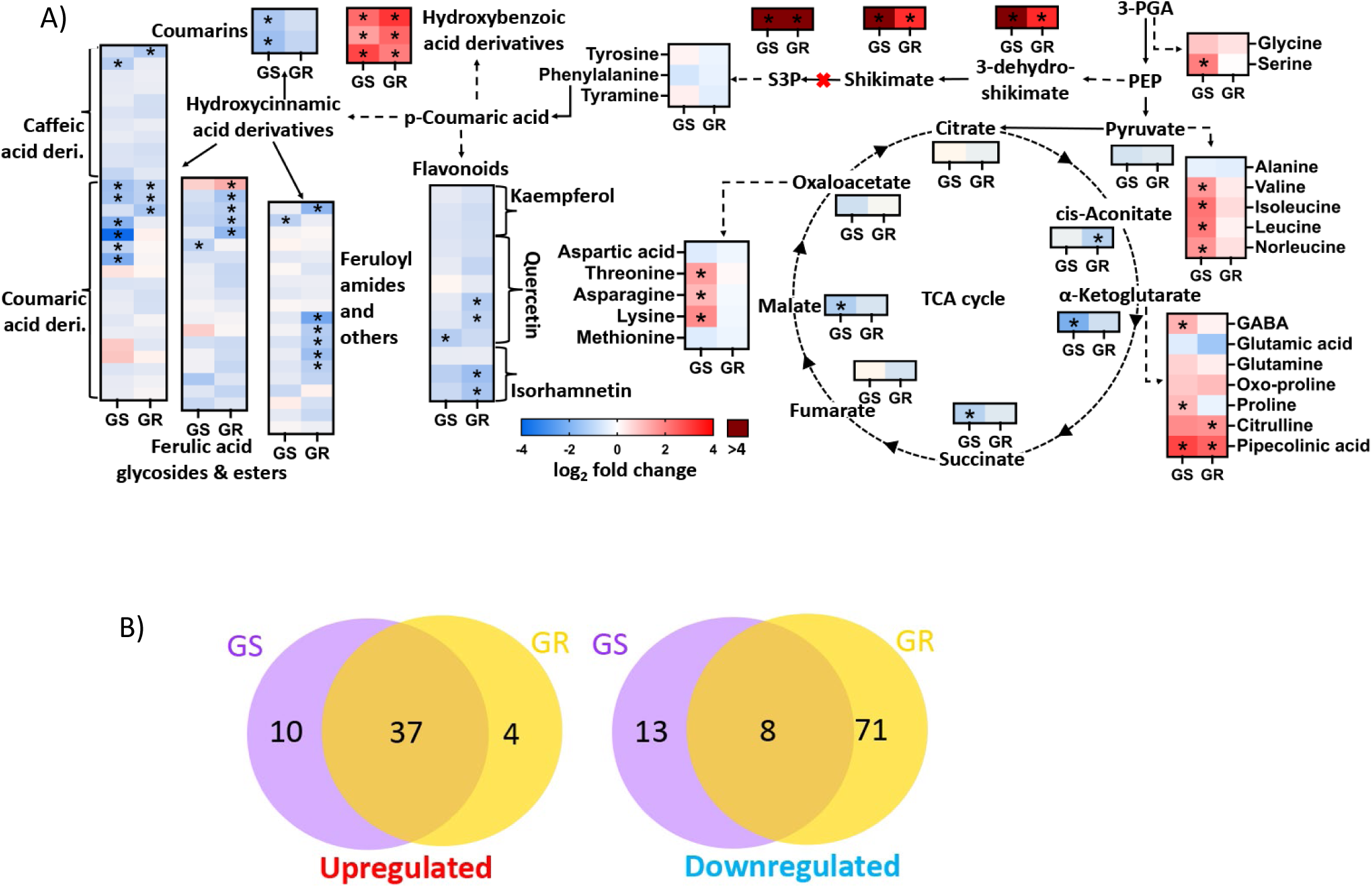
Outline of Tricarboxylic acid (TCA)-cycle metabolites, amino acids (primary) and identified phenylpropanoids outlined in the biosynthesis pathway (A) and Venn diagram showing the upregulated and downregulated secondary metabolites (B) in Glyphosate-susceptible (GS-) and resistant (GR-) biotype (glyphosate-treated vs control) of Amaranthus palmeri. Asterisk denotes significant difference (p<0.05, FDR<0.05, log2 fold change >|1|). Figure 7. Integrated pathway enrichment and network analysis of glyphosate-induced molecular changes in Palmer amaranth biotypes. Bar plots showing the top significantly perturbed pathways in glyphosate-susceptible (A) and glyphosate-resistant (B) Palmer amaranth biotypes, based on integrated analysis of differentially expressed genes, proteins, and metabolites using PaintOmics. Pathways are ranked by combined significance (p-value), with contributions from transcriptomic (green), proteomic (red), and metabolomic (blue) data, as well as the integrated (purple) score. KEGG network visualizations for GS (C) and GR (D) biotypes, illustrating the relationships among significantly impacted pathways and key metabolic hubs. Nodes represent pathways, and edges indicate shared molecular features or functional associations.

### Effect of glyphosate treatment on the aromatic amino acid biosynthesis pathway

Since EPSPS enzyme, the target of glyphosate, is a key driver of the shikimate pathway in plants (Maeda and Dudreva, 2012), we examined the glyphosate-induced changes in the transcript, protein, and metabolites of shikimate pathway. Forty transcripts, twenty-eight proteins, and five metabolites beginning from erythrose-4-phosphate and phosphoenolpyruvate to the biosynthesis of aromatic amino acids were detected [Figure 6; Figure S4]. In the absence of stress, the GR showed a ten times higher abundance of EPSPS transcript and protein compared to the GS [Figure S4] along with a higher abundance of shikimate and 3-dehydroshikimate. Additionally, GR in the non-stressed treatment had higher abundance of tyrosine aminotransferase transcript compared to the non-stressed GS while the non-stressed GS had a higher abundance of tryptophan biosynthesis transcript and proteins [Figure S4]. Upon glyphosate exposure, both biotypes showed an upregulation of transcripts of the sub-pathway that converts chorismate to tryptophan; however, the abundance of corresponding proteins was not affected [Figure S4]. Compared to the respective control, glyphosate-treated GS-also showed an upregulation of transcripts responsible for conversion of chorismate to phenylalanine/tyrosine that was not observed in the glyphosate-treated GR [Figure S4]. Protein abundance was mostly unaffected by glyphosate treatment except for the accumulation of anthranilate synthase beta subunit 1 across the biotypes and DAHP synthase 2 in GS [Figure S4]. In metabolites, the intermediates, including 3-dehydroshikimate, shikimate and shikimate-3-phosphate were accumulated across the biotypes – higher in GS than GR – while the final products, aromatic amino acids were not affected in abundance.

### Integration of transcriptomic, proteomic, and metabolomic data

To further explore the association between different levels of molecular activities in GR and GS with glyphosate treatment, we selected all the differentially expressed genes, proteins, and metabolites for integrated analysis in PaintOmics. As Palmer amaranth annotation is unavailable, we used the corresponding orthologous IDs of *Arabidopsis thaliana* in PaintOmics analysis. The transcriptomic, proteomic and metabolomic features were mapped to 142 pathways, with 24 pathways for GS and 13 pathways for GR showing significant perturbation (combined p ≤ 0.05) [Figure 7A and 7B; Table S12 and Table S13]. In GS, the Photosynthesis - antenna proteins (15 genes), Photosynthesis (39 genes, 1 metabolite) and Carbon fixation by Calvin cycle pathways (49 genes, 5 metabolites) shows a highly significant combined p-value of 2.32362E-15, 1.48714E-07 and 2.71339E-07, respectively, indicating severe disruption in light-harvesting complexes critical for energy production and carbon assimilation. This result is further supported by the KEGG network, which shows a higher impact in photosynthesis and secondary metabolism, and non-specific effect of glyphosate beyond the shikimate pathway [Figure 7C].

**Figure 7.**
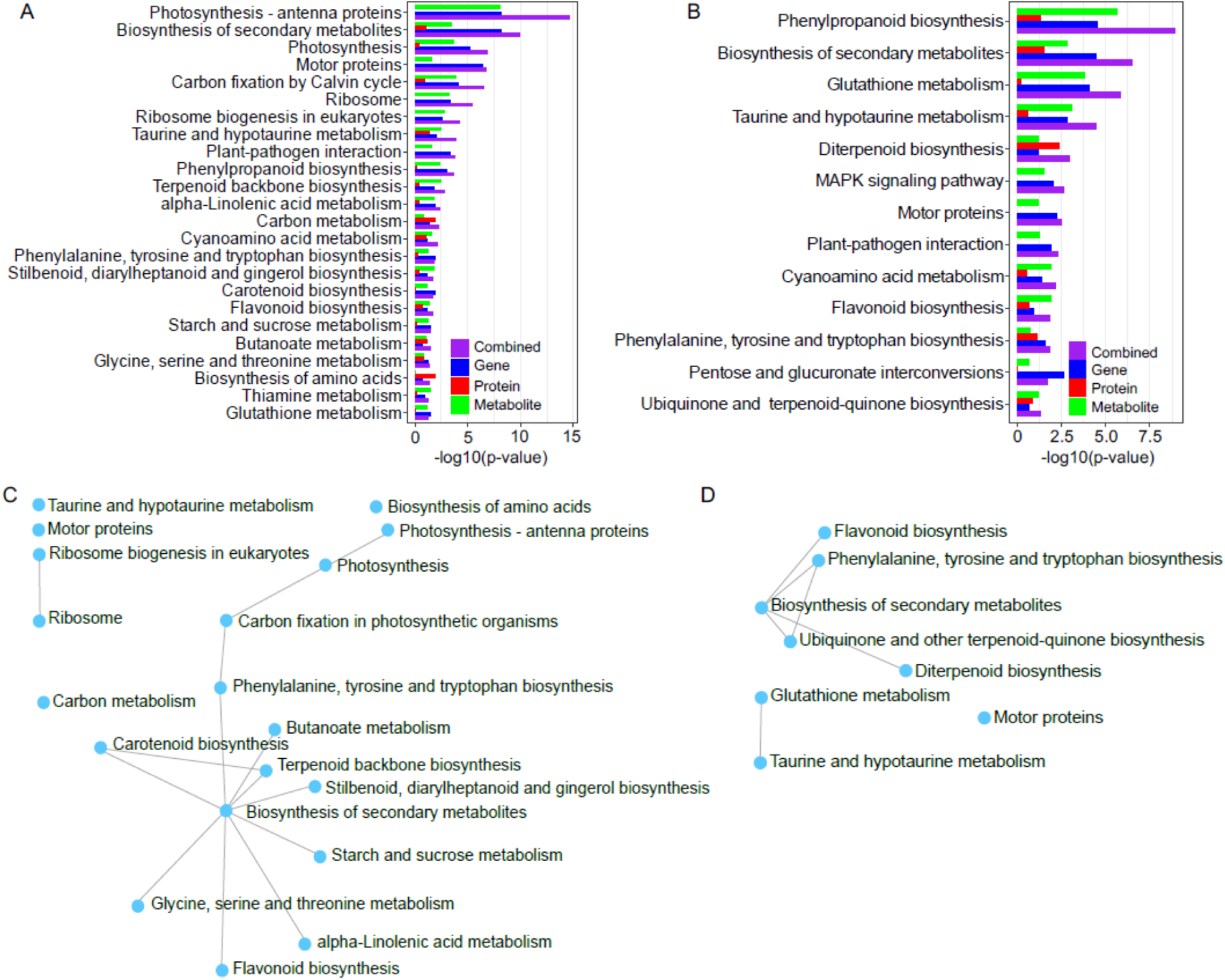
Integrated pathway enrichment and network analysis of glyphosate-induced molecular changes in Palmer amaranth biotypes. Bar plots showing the top significantly perturbed pathways in glyphosate-susceptible (A) and glyphosate-resistant (B) Palmer amaranth biotypes, based on integrated analysis of differentially expressed genes, proteins, and metabolites using PaintOmics. Pathways are ranked by combined significance (p-value), with contributions from transcriptomic (green), proteomic (red), and metabolomic (blue) data, as well as the integrated (purple) score. KEGG network visualizations for GS (C) and GR (D) biotypes, illustrating the relationships among significantly impacted pathways and key metabolic hubs. Nodes represent pathways, and edges indicate shared molecular features or functional associations.

The Biosynthesis of secondary metabolites pathway (736 genes, 92 metabolites) yielded a combined p-value of 1.19936E-10, suggesting perturbed defense pathways phenylpropanoids and terpenoids, essential for stress response. Pathways such as Taurine and hypotaurine metabolism (p-value 0.000140198) further indicate disrupted stress signaling and detoxification. Metabolic hub analysis identifying ferulic acid (31 DE neighbors, p-value 2.6481e-04) and L-alanine (6 DE neighbors, p-value 0.01572) as key hubs, underscoring the impact of glyphosate on amino acid and secondary metabolite networks via shikimate pathway inhibition. Pathways like Ribosome (p-value 3.80534E-06) and Motor proteins (p-value 1.70708E-07) suggest broader cellular impacts, affecting protein synthesis and transport, leading to cellular dysfunction in GS plants. Conversely, GR plants exhibit adaptive mechanisms to counter glyphosate as indicated by a high combined p-value 1.44911E-06 for Glutathione metabolism (46 genes, 6 metabolites), suggesting activation of detoxification mechanisms. The KEGG network for GR plants showed nodes around secondary metabolism and detoxification, suggesting resistance through shikimate pathway compensation and antioxidant defenses [Figure 7D]. Phenylpropanoid biosynthesis is more significant in GR (1.1553E-09) than GS (0.000238957), indicating less impact of glyphosate on shikimate pathway and downstream Biosynthesis of secondary metabolites.

Cellular pathways like Ribosome (GS: 3.80534E-06; GR: not significant) and Motor proteins (GS: 1.70708E-07; GR: 0.00311183) are more disrupted in GS, indicating broader cellular collapse, whereas GR maintains overall functionality.

## DISCUSSION

### Broad and coordinated molecular reprogramming induced by glyphosate highlights redundancy in plant stress-responses

Despite being a specific inhibitor of EPSPS (Duke and Powles, 2008), our multi-omics analysis shows that glyphosate caused extensive molecular reprogramming across multiple pathways, at the transcriptome, proteome, and metabolome levels in both GR and GS within 24 hours after treatment (HAT). This pervasive response likely arises from a combination of off-target toxicity of glyphosate (Campbell et al. 1976; Vaughn and Duke 1986; Kielak et al., 2011), and cascading consequences of the inhibition of the shikimate pathway as suggested by abundance of upstream metabolites including shikimate and 3-dehydroshikimate in both biotypes, despite an unchanged aromatic amino acids concentration at 24 HAT. In line with this commonality in responses, we observed activation of a core stress response network in both GS and GR. More than half of the transcripts induced by glyphosate were common to both GS-and GR-biotypes, highlighting substantial overlap in their early stress-response mechanisms. Several signaling and defense-related proteins were induced at transcript and protein levels across both biotypes 24 HAT, including kinases, a major regulatory protein in plant response to stressors (Faus et al, 2015; Kim et al., 2017; Wang et al., 2020) that accounted for ∼15% of the upregulated transcripts.

Transcriptional and translational upregulation of pathogenesis-related (PR) proteins in the ABA-mediated signaling pathway, calmodulins, calcium-dependent kinases, and other pathogen-induced genes and proteins that are less directly attributable to inhibition of the shikimate pathway were also observed 24 HAT. These changes reflect the induction of a generalized stress signaling cascade that is promoted by the ability of glyphosate to increase endogenous levels of ABA and gene expression of ABA-mediated signaling pathway (Fuchs et al., 2021) observed in *Cyperus esculentus* (Cañal et al., 1990) and *Glycine max* (Jiang et al., 2013), respectively. The role of ABA-mediated signaling, coupled with the involvement of key regulatory proteins such as kinases and PR, illustrates the complexity and redundancy of plant defense systems. The upregulation of multiple stress-response pathways emphasizes the interconnected nature of plant defense, where a herbicide that is thought to inhibit a single enzyme and cause a slower plant death can trigger a multifaceted defense system integrating both abiotic and biotic signaling networks within 24 HAT. This rapid, broad-spectrum response brought about by a unified signaling network likely provides a survival advantage through redundancy in stress response.

Beyond ABA-mediated signaling, our data suggest that glyphosate also perturbs jasmonic acid (JA) associated pathways. Differential expression of several JA biosynthesis genes, such as lipoxygenases, allene oxide synthase, and oxophytodienoate reductase, were observed following glyphosate treatment, indicating that multiple phytohormone networks are engaged in the herbicide response (Fuchs et al., 2021. The involvement of both ABA and JA, alongside downstream defense responses including PR proteins, underscores the breadth of the induced signaling cascade. Furthermore, the chitin catabolic activity mediated by endochitinases, a typical pathogen-related defense mechanism, was upregulated specifically in the GR (Kumar et al., 2018; Zhang et al., 2014). The induction of chitinase, together with calcium-dependent signaling, suggests that glyphosate stress can trigger biotic defense components in resistant plants, potentially reflecting an enhanced defensive capacity or a form of stress priming in GR.

### Glyphosate induces contrasting perturbation of primary metabolism in GS and GR biotypes

In contrast to their shared inducible defense, the two biotypes showed contrasting patterns in perturbation of primary metabolism, with a severe inhibition of central carbon metabolism occurring only in GS. Integrated pathway analysis revealed that in GS the photosynthesis light harvesting complex, and carbon fixation pathways were among the most significantly perturbed. Multiple components of central carbon metabolisms, including photosystems, the Calvin cycle, energy cycling, amino acid, and lipid biosynthesis, were transcriptionally and translationally downregulated in GS, which in turn contributes to the observed disruption in carbon assimilation and energy generation. Such glyphosate-induced dysregulation of central carbon metabolism (Orcaray et al., 2012; Sandhu et al, 2023) and photosynthesis (Gomes et al.,2016a; 2016b; 2017) has been noted at the metabolite level in prior studies and could contribute to off-target toxicity underlying plant death (Gomes et al., 2014). The GR was able to circumvent this glyphosate-induced system-wide perturbation, which indicates a protection from metabolic disruption. Even though, compared to the GS, the baseline gene expression and protein abundance of the target enzyme-EPSPS was 10 times higher in the GR than in GS, glyphosate did not induce any further changes in EPSPS expression at 24 HAT in either biotype. This indicates that GS injury is not due to differential EPSPS regulation, but rather downstream consequences of its inhibition, or off-target toxicity. We speculate that the known genetic adaptation of GR, such as the co-amplification of stress-related genes (Koo et al., 2018), buffer the GR from the physiological impact of the herbicide, allowing GR to preserve primary metabolic flux.

Our network analysis supports this notion, where the key hub metabolites such as ferulic acid and L-alanine were disrupted only in GS, indicating that the GS experienced broader metabolic perturbation linking central carbon and secondary metabolism that the GR was able to circumvent. The integrated pathway analysis revealed that the broad secondary metabolites were more impacted in the glyphosate-treated GS than in the GR. This corresponds to a marked suppression of phenylpropanoid and terpenoid biosynthesis in the susceptible plants, depriving them of many defense-related secondary compounds. The collapse of these protective secondary metabolic pathways in GS likely compounds the effects of primary metabolic perturbation, making GS plants highly vulnerable to oxidative damage. In contrast, the GR showed no such collapse of secondary metabolism, and potentially rerouted metabolic flux toward phenylpropanoids to bolster detoxification. Thus, glyphosate-treated GS suffer a combined breakdown of both primary and secondary metabolism, whereas GR maintain core metabolic functions and defensive metabolite production under herbicide stress.

### Integrative multi-omics analysis reveals distinct response mechanisms in GS and GR biotypes

Unlike the GS, the GR exhibited a targeted detoxification and antioxidant defense response. While the glutathione metabolism was not identified as the top changing pathway in any of the single omics data sets, the integrated multi-omics analysis revealed that this pathway was the most significantly upregulated in GR. No significant changes were detected in the metabolite levels of oxidized glutathione (GSSG) in either biotype (Supplementary Table S12), suggesting that the glutathione pathway activation is evidenced mainly at the transcript/protein level. This suggests that multi-layer data integration is required to capture complex plant responses with high confidence, as evident from the increased statistical significance. Previous studies from our lab, based on enzyme functional activity, identified the upregulation of the antioxidant machinery in GR upon exposure to glyphosate (Maroli et al. 2015), which supports the results from the current data integration analysis, and highlights the ability of the omics integration to inform functional outcomes. Glutathione plays a crucial role in mitigating herbicide stress by conjugating glyphosate or neutralizing oxidative damage, and our data indicate that GR uniquely activates this mechanism. Similarly, the phenylpropanoid pathway exhibited a higher perturbation in GR than in GS, which suggests an active rerouting of metabolic flux towards the production and utilization of compounds with a higher antioxidant potential. Although transcriptomic data indicated an up-regulation of phenylpropanoid biosynthetic genes and flavonoid biosynthesis in GR response to glyphosate, compared to the native level, many phenylpropanoid metabolites, including caffeic and ferulic acid conjugates, quercetin glycosides, significantly diminished in GR plants after treatment (Supplementary Table S12). This suggests that GR was actively using or remodeling these antioxidant compounds during glyphosate stress. Thus, the upregulation of phenylpropanoids is evidenced by gene/pathway activation, even though the outcome is a net decrease in some phenolic metabolites, potentially due to their consumption in ROS scavenging or other protective processes. This interpretation is supported by the KEGG network analysis, which showed that the perturbed nodes in GR were clustered around secondary metabolism and detoxification processes, in contrast to the GS network, which was dominated by photosynthetic and other primary metabolic nodes, and suppression of phenylpropanoid biosynthesis. The GR also largely maintained the integrity of core cellular functions, as observed by the lack of downregulation of ribosomal proteins and motor proteins that are important for protein synthesis and intracellular transport, whereas they were strongly repressed in GS. The genetic basis of this resilience includes EPSPS gene amplification and the presence of extrachromosomal circular DNA carrying defense-related genes (Gaines et al., 2010; Koo et al., 2018). The system-level adaptation that upregulates detoxification pathways and sustains vital processes in GR illustrates the complexity of glyphosate resistance beyond the canonical target-site mechanism.

Taurine and hypotaurine metabolism is another instance where metabolic disruptions were evident only upon integrative analysis, especially since this specific taurine biosynthesis pathway is less reported in plants, and hence, the pathway-impact would be more driven by the associated metabolites (Lähdesmäki 1986). Although modest changes in related genes, proteins, or metabolites alone did not meet individual significance cutoffs, the combined multi-omics data identified this pathway as significantly perturbed under glyphosate stress in both biotypes.

Taurine is a sulfur-containing non-proteinogenic amino acid derivative known to confer osmoprotection and antioxidant protection (Farooq et al., 2025). Perturbation of this pathway, particularly pronounced in the GS plants, indicates oxidative or osmotic stress imposed by glyphosate. A greater osmotic stress in glyphosate-treated GS is also supported by a 2-fold increase in proline content (Supplementary Table S11), whereas this osmolyte remained unchanged in GR following the herbicide exposure. Likewise, our integrative analysis pointed to alterations in carbohydrate metabolism related to raffinose family oligosaccharides (RFO) in the GR, a class of sugars known to accumulate as osmoprotectants and ROS-scavengers during plant stress (Nishizawa et al., 2008. While subtle, such changes hint that the resistant plants may reallocate carbon resources toward protective osmolytes as part of their defense strategy, possibly by up-regulating RFO biosynthesis genes and diverting sugars into those pathways, a characteristic that evaded single-layer analyses. For example, raffinose itself was not identified as a significantly changed metabolite in the metabolomics dataset, and genes of the RFO biosynthetic pathway, such as galactinol synthases, were not prominent in the standalone transcriptomic analysis. Because RFOs act as both osmoprotectants and ROS-scavengers, their coordinated biosynthetic activation complements the glutathione-and phenylpropanoid-based defenses that were also enhanced in GR plants. These complex responses emerged primarily through integrating omics layers, reinforcing that multi-omics integration could be an ideal approach to uncover the stress response mechanism in plants.

### The strong correspondence between transcript and protein abundance suggests tight regulation of transcription and translation in Palmer amaranth

The multi-omics data also provide insight into the regulatory coordination between gene expression and protein synthesis under glyphosate stress. Overall, a strong positive association (r > 0.50) was observed between the transcripts and protein abundance under glyphosate stress, similar to that reported in *A. thaliana* (Mergner et al. 2020). These results contrast with earlier studies that reported a poor coregulation of transcriptome and proteome (De Godoy et al., 2008; Fournier et al., 2010; Srivastava et al., 2013; Bathke et al., 2019). At the metabolite level, transcript changes aligned better with shifts in primary metabolite levels than with protein changes, whereas the correlation with secondary metabolites was poor. This suggests that many primary metabolites are transcriptionally regulated, while protein abundance alone is not a direct predictor of metabolite accumulation, potentially due to post-translational control (de Visser et al. 1992; Ishihara et al., 2017). Despite the general transcript-protein coordination observed, the GS exhibited potential translational inhibition under glyphosate stress that was not present in GR. Specifically, the integrated analysis identified significant downregulation of ribosomal proteins and biogenesis factors in GS, implying that protein synthesis capacity was hampered in the susceptible plants. Consistent with this, a comparison of protein-to-transcript ratios (PTR) in each treatment (Figure S3) showed a reduction in the median PTR in glyphosate-treated GS relative to untreated controls, whereas the GR biotype maintained higher PTR values. This could be interpreted as GS producing fewer proteins per mRNA under herbicide stress, a likely consequence of energy deficits and damage to the translation machinery, which is partly supported by the marginal decrease in total soluble proteins in leaves of GS treated with glyphosate (Figure S6). This translational disruption was not observed in GR, as it maintained robust ribosomal function and energy status. This difference further highlights how GS succumbs to broad physiological disruption, while GR sustains growth processes under glyphosate stress.

Overall, our multi-omics approach reveals that glyphosate imposes a complex stress syndrome involving far-reaching changes beyond its primary enzymatic target. GS activate a generalized defense response but undergo extensive metabolic disruption, resulting in failure of energy production and biosynthetic systems, which likely accelerates plant death. In contrast, GR activates a coordinated set of protective pathways and sustains less disruption of primary metabolism, enabling them to withstand the herbicidal stress with minimal physiological compromise. By combining transcriptomic, proteomic, and metabolomic data, we uncovered the herbicidal impact on critical pathways that would have otherwise remained less obvious. This comprehensive perspective provided not only a list of perturbed pathways but also a deeper understanding of the regulatory networks and compensatory mechanisms at play across multiple levels of cellular function. Such insights are invaluable for advancing plant stress physiology research.

## MATERIALS AND METHODS

### Plant growth and treatment application

Seeds from GR biotype, T4B1 (GR_50_: 1.3 kg ae ha^-1^, EPSPS gene copy number >33), and GS biotype, S17 (GR_50_: 0.09 kg ha^-1^, EPSPS gene copy number: 1) (Nandula et al., 2012, Sandhu et al, 2023), of Palmer amaranth were sourced from the Crop Production Systems Research Unit in Mississippi, US (USDA-ARS, Stoneville, MS). Plants were grown in a greenhouse, supplemented with metal halide lamps for the duration of the experiment. Parent plants were raised from seeds sown in 48-cell trays in the germination mix (SunGro). After two weeks of growth, seedlings were transplanted to germination mix in pots (15 cm diameter x 11 cm depth) and were fertilized by soil application of 50 mL of 4 g L^-1^ MiracleGro solution (MiracleGro, 24%−8%−16% [N−P−K], Scotts Miracle-Gro Products, Inc., Marysville, OH) one week after transplanting. Plants of both biotypes were obtained by propagating axillary buds of parent plants as described in (Hoagland et al., 2013; Ribeiro et., 2014). Twelve days after propagation, plants were transplanted to germination mix in pots (10 cm diameter x 9 cm depth) and were fertilized and watered as needed. The plant growth conditions, and plant care have been optimized for these biotypes and detailed in previous studies (Nandula et al., 2012; Maroli et al., 2015, 2017; Sandhu et al., 2023).

Twelve days after transplanting (27 days after propagation), five plants per biotype were sprayed with 0.42 kg a.e. ha^-1^ of glyphosate (Roundup ProMax, Monsanto Co, St. Louis, Mo), which is half the field recommended dose, using a Teejet 8001EVS nozzle in an enclosed spray chamber (DeVries Manufacturing, Hollandale, MN). Five non-sprayed plants served as untreated controls in each biotype. Twenty-four hours after treatment (HAT), the actively growing top whorl of leaves along with the top meristem were harvested from glyphosate and control treatments and stored at-80°C until analysis following the previously optimized protocols (Maroli et al., 2015, 2017, Sandhu et al. 2023). Prior to analysis, the leaf tissue was transferred to 50 mL centrifuge tubes and ground using Geno/Grinder® (SPEX SamplePrep, Metuchen, NJ, USA). The tissue was ground for three cycles of 1,750 rpm for 45 seconds each after the addition of 15 metallic beads. The tubes were kept frozen throughout the grinding using liquid nitrogen to avoid sample degradation. The ground samples were stored at-80°C and used for further analysis.

### Metabolite extraction

The polar metabolites were extracted as per Sandhu et al. (2023) using 80% methanol (w/v) in a ratio of 1:10. Briefly, 80% methanol (w/v) was added to 100 mg of ground tissue. Samples were homogenized with ceramic beads at 6,000 rpm for 30 seconds for eight cycles at 1°C. The homogenized samples were centrifuged for two cycles at 13,500 rpm for two minutes at 0°C. 200 µL of methanol extract was transferred to new tubes followed by the addition of ice-cold chloroform and water in a ratio of 1:1. After centrifugation at 13,500 rpm for one minute at room temperature, the top methanol-water phase was removed and used for downstream primary and secondary metabolomic analysis.

### Untargeted primary metabolomics analysis by GC/Q-ToF

Chloroform partitioned methanol-water phase was used to quantify primary metabolites using Gas chromatography/quadrupole-time-of-flight mass-spectrometry (GC/Q-ToF). Briefly, 20 µL of the methanol-water extract was dried down in a glass vial with 10 µL of internal standard mix containing 20 ppm ribitol and 50 ppm d27-myristic acid in hexane (Sandhu et al., 2023). The dried samples were methoxylaminated at 40°C with 20 µL of methoxylamine hydrochloride (40 mg mL^−1^) in pyridine for 90 min followed by silylation of the metabolites with 100 µL of N-methyl-N-(trimethylsilyl) trifluoroacetamide (MSTFA) with 1% trimethylchlorosilane (TMCS) and 1 ppm alkane mixture (C27-C30) for 40 minutes at 40°C. A method blank and three pooled samples were also derivatized for quality control (QC). A serial dilution of 52 compound standard mix (1, 2, 3, 5, 8, 10, 20, 30, and 40 ppm) were also derivatized as above to aid in compound identification. Trimethylsilyl derivatives of the metabolites were separated on a DB-5MS UI column (30 m x 0.25 mm, 0.25 film, Agilent Technologies, Santa Clara, CA, USA) using gas chromatography (GC) (Agilent 7890B GC) and analyzed on a quadrupole time-of-flight mass spectrometer (Agilent 7250 GC/Q-ToF). One microliter of the sample was injected into the GC inlet kept at 280°C. Helium was used as the carrier gas with the flow rate set to a constant pressure of 9.6 psi and a split ratio of 25:1. The column oven temperature gradient was as follows: initial hold at 60°C for 1 minute, followed by a linear ramp of 10°C min^-1^ to 310°C, and a hold at 310°C for 10 minutes. The source inlet temperature was set to 280°C, the electron ionization (EI) voltage was set at 70 eV and scan range of m/z 60 – 460. Mass spectra were processed using MS-Dial (Tsugawa et al., 2011a; 2011b; Matsuo et al., 2017). The same weightage was given to retention time (RT) and EI similarity. The identification of the primary metabolites was based on Kovats retention index (RI), and the mass fragmentation pattern matches with the standard mix and Kovats RI Library (GCMS DB_AllPublic-KovatsRI-VS2). The metabolite confirmed with authentic standards was identified at level 1 of the Metabolomics Standards Initiative (MSI), while those matching the library were annotated at level 2 of MSI. The minimum EI similarity score cut off was set to 75% with an RI tolerance of 20. RT was used for alignment with RT tolerance of 0.075 min, EI similarity tolerance 75%, RT factor 0.5, EI similarity factor 0.5.

### Untargeted metabolomics analysis by UHPLC-MS/MS

The top methanol-water phase was also analyzed on ultra-high performance liquid chromatography (UHPLC) coupled to an Orbitrap Fusion Tribrid mass spectrometer (Thermo Scientific, Waltham, MA) to capture metabolites that are less amenable to GC analysis. 150 µL of isotope-labeled resveratrol (^13^C_6_; 2 ppm; internal standard) was added to 150 µL of the sample. Quality control (QC) samples were prepared by pooling equal aliquots from all individual samples and run after every 12 samples. The liquid chromatography separation was done on an Dionex UltiMate 3000 UHPLC system (Thermo Fisher Scientific). A sample aliquot of 5 µL was injected onto an Acquity UPLC HSS T3 column (2.1 × 150 mm, 1.8 µm, 100 Å; Waters, Milford, MA, USA) held at 32 °C. Mobile phase A was water containing 0.05 % formic acid and mobile phase B was neat acetonitrile containing 0.05 % formic acid with a solvent flow rate of 0.20 mL min-1. The gradient proceeded from an initial 5 % to 60 % B over 15 min, followed by a 2 min wash at 90 % B and a 5 min re-equilibration at 5 % B. Ionization was performed with a heated-ESI (HESI-II) source at 3.5 kV (positive mode) or 3.0 kV (negative mode). Sheath, auxiliary and sweep gases were set to 15, 5 and 1.5 mL min-1, respectively. The vaporizer and ion-transfer-tube temperatures were held at 300 °C. Full-scan MS data were acquired in an Orbitrap at 60,000 FWHM (m/z 200) over m/z 120–1200 with an AGC target of 4 × 105 and a 50 ms maximum injection time. Data-dependent MS/MS used monoisotopic precursor selection, an intensity threshold of 2 × 105, a 10 s dynamic exclusion (±10 ppm) and HCD at 40 % Normalized Collion Energy. Product ions were recorded at 15,000 FWHM with an AGC target of 5 × 104 and a 22 ms injection time.

Raw files from positive and negative ionization were processed separately in Compound Discoverer 3.1 (Thermo Fisher Scientific). Features eluting < 2.0 min, > 17.0 min, or with a peak area < 1 × 10 6 were discarded. Positive-mode data provided 5016 mass features and served as the primary dataset; peaks unique to, or much more abundant in, negative mode (2904 mass features) were taken from the negative file. After consolidation of the adducts, in-source fragments and exclusion of peaks with low signal-to-noise and asymmetry, mass features were annotated by accurate mass (< 5 ppm), retention time and MS/MS fragmentation against an in-house library (Sandhu et al., 2023), MoNA, mzCloud, Flavonoid Search, HMDB and published literature. Resulting identifications were assigned to MSI levels 2 or 3. The data from UHPLC-MS/MS was also processed on SIRIUS v.4.4 (Dührkop et al., 2019) to predict the ClassyFire superclass and class (Djoumbou Feunang et al., 2016) of unidentified compounds using CANOPUS (Dührkop et al., 2021). The raw files were converted to mzML and imported directly into SIRIUS. The features were scored for MS/MS isotopes (Böcker et al., 2009) with an MS^2^ mass deviation of 5 ppm. The elemental composition was limited to molecular formulas available in databases. [M+Na]+, [M+K]+ and [M+H]+ adducts were allowed for molecular formula prediction while [M+NH4]+ and [M+ACN+H]+ were additionally included as fallback adducts for CSI:Finger ID (Dührkop et al., 2015; Böcker and Dührkop, 2016; Hoffmann et al., 2021). ZODIAC (Ludwig et al., 2020) was also used to improve the annotation of molecular formulas.

### Protein extraction and digestion

Proteins were extracted from leaf samples (∼100 mg frozen leaf) with lysis buffer normalized to a ratio of 1:10 leaf to lysis buffer (w/v). The lysis buffer consisted of 0.1 M Triethylammonium bicarbonate buffer, 5% (w/v) sodium dodecyl sulfate, 0.023% (v/v) phosphoric acid, 1.185X Halt™ protease inhibitor, and 0.56% Tris(2-carboxyethyl)phosphine hydrochloride (TCEP) (w/v) in water. Samples were homogenized with ceramic beads at 6,000 rpm for 30 seconds and sonicated at 25% amplitude for 50 seconds and centrifuged at 12,000 rpm for 2 minutes. For protein reduction with TCEP, 400 μL of supernatant was transferred to 600 μL Eppendorf tubes and incubated at 95°C for 15 minutes. Reduced proteins were alkylated by adding 50 μL of 400 mM iodoacetamide to samples and incubating in dark at room temperature for 30 minutes.

Soluble proteins in the extracts were quantified using Bicinchoninic Acid-Reducing Agent Compatible (BCA-RAC) assay as specified in the manufacturer’s protocol (Thermo Scientific) and 150 μg of protein loaded to S-trap mini spin columns following the manufacturer’s protocol (Protifi, Farmingdale, NY). Proteins were digested using trypsin (21:1, protein to trypsin, w/w) containing 0.01% ProteaseMAX surfactant and incubated for 16-hours in a water bath maintained at 37°C. Peptides were eluted from S-trap mini spin columns following the manufacturer’s protocol (Protifi), dried *in vacuo*, and dissolved in 150 μL 0.1% formic acid in water (equivalent to 1 μg/μL protein). Samples were diluted by adding 5 μL 0.5 pmol/μL Pierce Peptide Retention Time Calibration Mixture (PRTC) as an internal standard to 45 μL samples.

### Proteomics using NanoSpray LC-MS/MS

Proteins (1 μg load) were separated on an EASY-Spray PepMap RSLC C18 column (75 μm x 50 cm, 2 μm, 100 Å) using an UltiMate 3000 RSLCnano (Thermo Scientific) liquid chromatograph coupled to an Orbitrap Fusion Tribrid mass spectrometer (Thermo Scientific) equipped with an EASY-Spray ion source (Thermo Scientific). The mobile phases A and B consisted of 0.1 % formic acid in water and 0.1% formic acid in 80% acetonitrile, respectively. The solvent gradient began with a three-minute hold at 4% B, followed by a three-step ramp to 30% B at 90 minutes, to 50% B at 115 minutes, to 90% B at 125 minutes, holding at 90% B for 20 minutes and then re-equilibrated at 4% B for 30 minutes. A flow rate of 0.15 μL/min and injection volume of one μL was used. Peptides were ionized in an EASY-Spray ion source (Thermo Scientific) operated in positive ionization and peptide mode with a default charge state of 2, spray voltage of 2.2 kV, sweep gas 3 L/min, and ion transfer tube temperatures of 275°C. Precursor ions were analyzed using Orbitrap with a resolution of 50 K FWHM, the scan range of m/z 375-1500, AGC target of 4e5, and maximum injection time of 50 ms. Filters for peptide MIPS, intensity threshold of 1.9e4, charge state 2-7, and dynamic exclusion duration of 40 seconds with 10 ppm mass tolerance. DDA MS^2^ fragmentation was performed using collision-induced dissociation (CID) of 35% and product ions were analyzed in Ion Trap with a rapid scan rate. For processing, the acquired mass spectra were processed in Proteome Discoverer v.2.2 (Thermo Scientific) using Label-Free Quantification workflow as per manufacturer instructions. Mass spectra were aligned to peptides that would be predicted from the FASTA file created from GR Palmer amaranth female plant using Sequest HT algorithm. The processing workflow in Proteome Discoverer included Spectrum Files RC, Spectrum selector, Sequest HT, Percolator, and Minora feature detector nodes. Default parameters were used except for a few modifications. Carbamidomethyl (+57.021 Da (C)) was added as a static modification for the Spectrum Files RC. In the Sequest HT node, precursor mass tolerance against the database was set to 10 ppm with a fragment mass tolerance of 0.6 Da, and oxidation (+15.995 Da (M)) was added as a dynamic modification while acetyl (+42.011 Da (N)) was added as N-Terminal modification. The MSF files created from the processing step were used in the consensus workflow for label-free quantification. The consensus workflow nodes were PSM grouper, peptide validator, peptide and protein filter, protein scorer, protein grouping, peptide in protein annotation, protein FDR validator, protein marker, feature mapper, and precursor ions quantifier. The precursor quantification was based on intensity, and the abundance was normalized based on total peptide amount without any scaling. Missing values were imputed based on low-abundance resampling.

### Statistical analysis of untargeted primary and secondary metabolomics and proteomics

The statistically significant differences between the treatments for untargeted primary and secondary metabolomics and proteomics data were obtained by performing the pairwise comparisons (parametric t-test) between the glyphosate and control treatments within GS and GR in Graph Pad Prism v9. Multiple comparisons were controlled by the False Discovery Rate (FDR) approach using a two-stage step-up method of Benjamini, Krieger, and Yekutieli (Benjamini et al., 2006). Proteins/metabolites with p<0.05, FDR <0.05 and log_2_ fold change>|1| were considered significantly different between treatments.

### RNA extraction, sequencing and analysis

Total RNA was extracted using TRIzol® Reagent (Invitrogen, Thermo Fisher Scientific, USA) as per manufacturer instructions. Briefly, 1 mL of cold TRIzol® Reagent was added to 100 mg of finely ground tissue and vortexed vigorously for proper mixing. The lysate was centrifuged at 12,000 g for 5 minutes at 4°C and the clear supernatant was transferred to a new 2 mL centrifuge tube without disrupting the pellet. This was followed by incubation at room temperature for 5 minutes to permit the complete dissociation of nucleoprotein complexes. 200 μL of chloroform was added to the lysate, mixed properly, and incubated again for 3 minutes at room temperature.

After incubation, the samples were centrifuged at 12,000 g for 15 minutes at 4°C. The mixture separated into a lower red phenol-chloroform phase, interphase, and a colorless upper aqueous phase. 400-500 μl of the upper aqueous phase was transferred to a new tube without touching the interphase to avoid contamination from DNA and proteins. The aqueous phase was again partitioned with equal amounts of chloroform (1:1) to remove any inadvertent contamination.

The water:chloroform mixture was incubated on ice for 5 minutes followed by centrifugation at 12,000 g for 10 minutes at 4°C leading to phase separation. The top-aqueous phase was transferred to a new tube. At this step, total RNA was isolated by the addition of 0.6X of cold isopropanol. The solutions were mixed well by inverting the tubes approximately 30 times followed by incubation on ice for 5 minutes. For precipitation, the tubes were centrifuged at 12,000 g for 10 minutes at 4°C. Total RNA formed a white gel-like pellet at the bottom. The supernatant was discarded, and the RNA was washed with 1 mL of 75% ethanol for 1 hour at room temperature. Finally, the tubes were centrifuged (7,500 g for 5 minutes at 4°C) and the ethanol was removed completely using a micro pipettor and the pellet was airdried for 5 minutes. The dried total RNA pellet was dissolved by pipetting up and down in the 40 μL of DNase treatment solution (36 μL RNase free water + 4 μL of DNase buffer 1X + 1.5 μL DNase I). The integrity of RNA was determined by agarose gel electrophoresis (2:1 ratio of 28S:18S RNA) [Figure S5]. 150 bp paired-end mRNA sequencing (mRNA-seq) was performed by Novogene Corporation (Sacramento, CA) using illumina NovaSeq platforms. Prior to library construction, the RNA was analyzed on Bioanalyzer for quality control. The RNA integrity number of all the samples was 7 or above except one which was 6.9 [Figure S6]. Samples of 1–2 μg mRNA were used for library construction and sequencing. mRNA was purified from total RNA using poly-T oligo-attached magnetic beads. After fragmentation, the cDNA was synthesized followed by end repair, A-tailing, adapter ligation, size selection, amplification, and purification. Raw reads were filtered to remove adapters and low-quality reads (Qscore <=5 or undetermined bases more than 10%) [Table S1]. The mRNAseq data was assembled based on a modified procedure from Trinity using the alignment-based RSEM method. Briefly, raw FASTQ files were preprocessed to remove adapter and low-quality bases (Phred threshold = 33) with Trimmomatic software.

Cleaned reads were mapped to the Palmer amaranth genome sequenced from a GR female plant using the Bowtie2 short-read aligner. Transcript abundance was quantified with RSEM-1.3.3.

The abundance was normalized using the Trimmed Mean of M-values (TMM) method and expressed in Transcripts per million (TPM). The differential expression analysis was performed using edgeR. Transcripts with p<0.05, FDR <0.05 and log_2_ fold change>|1| were considered significantly different between treatments.

### Functional analysis

The differentially expressed transcripts and proteins (p<0.05, FDR <0.05 and log_2_ fold change>|1|) were analyzed for Gene Ontology (GO) enrichment analysis for functional interpretation. GO enrichment was performed on R v4.0 using Goseq v1.46 (Young et al., 2010). For pathway enrichment analysis, Palmer amaranth FASTA was blasted to find orthologs from *A. thaliana*. Orthologs with at least 40% similarity and the ones that were already identified from *A. thaliana* were retained.

### Correlation and protein-mRNA Ratio (PTR) analysis

Pearson correlation analysis between omics was performed between the log(x+1,2) transformed abundance values of proteins and transcripts. Protein to mRNA ratio (PTR) was calculated by dividing the abundance of protein with the corresponding transcript followed by log_2_ transformation as described by Mergner et al. (2020). PTR density distribution curves were made in R v4.0 using package ggplot2. Statistical analysis for PTR was also performed by pairwise comparisons (parametric t-test) and multiple comparisons were controlled using two-stage step-up FDR method of Benjamini, Krieger, and Yekutieli (Benjamini et al., 2006)

### Integrated analysis using PaintOmics

The transcriptomic, proteomic and metabolomic data was integrated using PaintOmics 4 (Liu et al., 2022). Differentially expressed genes, proteins, and metabolites identified from transcriptomic, proteomic, and metabolomic datasets were selected for integration. Due to the lack of comprehensive Palmer amaranth genome annotation, we mapped all features to their corresponding Arabidopsis thaliana orthologs using BLAST. The *Arabidopsis thaliana* reference organism was selected in PaintOmics for pathway mapping and enrichment analysis. The lists of differentially expressed genes, proteins, and metabolites (with their Arabidopsis ortholog IDs and metabolite names) were uploaded to PaintOmics and visualize their distribution across KEGG pathways. Pathways showing significant perturbation were identified based on combined enrichment statistics provided by PaintOmics, and results were further explored using the interactive network and pathway visualization tools within the platform.

## Supplementary data

**Figure S1. PLS-DA plot of first two components from PLS-DA of transcriptomics, proteomics, primary (GC-QToF), and secondary metabolomics (UHPLC-MS/MS) data.** Different treatments are represented by different colors and ellipses represent 95% confidence interval.

**Figure S2. PTR Density plot of the protein-transcript pairs.** Density plot of protein-mRNA ratio (PTR) for the protein with corresponding transcript abundance in control and glyphosate-treated GS and GR biotype of *Amaranthus palmer*i. The straight vertical line represents the median values for each treatment.

**Figure S3. Identified terpenoids of the present study.** Asterisk denotes significant difference (p<0.05, FDR<0.05, log2 fold change >|1|).

**Figure S4: Effect of glyphosate on transcripts and proteins of aromatic amino acid biosynthesis pathway in GS and GR *Amaranthus palmeri*.** Heatmap displaying the log_2_ fold change in the abundance of transcripts and proteins of aromatic amino acid biosynthesis pathway. Column C is the log_2_ fold change of GR control vs GS control. GS and GR respectively denote the log_2_ fold change of glyphosate treatment with respective control for GS and GR biotypes. First three columns are for transcript abundance and the next three for protein abundance. Asterisk denotes significant differences at p<0.05, FDR <0.05, and log_2_ fold change >|1|.

**Figure S5. Agarose gel electrophoresis of isolated total RNA from leaves of *Amaranthus palmeri*.** RNA bands (28S, 18S, 5S) are indicated by arrows. Samples with a ratio of 2:1 for 28S:18S bands are considered good quality. The ID on top of each gel column represents sample ID. For details about samples, refer to table S1.

**Figure S6. Total soluble proteins in leaves of *Amaranthus palmeri*.**

**Table S1.** Quality attributes of RNAseq analysis. Raw reads, error percentage, Phred scores (Q20, Q30), and GC percentage in each sample, including control (Ctrl) and glyphosate (Gly) - treated glyphosate-resistant (GR) and-susceptible (GS) biotypes of *Amaranthus palmeri*.

**Table S2**. Metabolites identified in control and glyphosate-treated GR-and GS-biotype of *Amaranthus palmeri* from GC-QToF analysis

**Table S3.** Metabolites identified in control and glyphosate-treated GR-and GS-biotype of *Amaranthus palmeri* from UHPLC-MS/MS analysis

**Table S4.** Classification of unidentified compounds from UHPLC-MS/MS analysis. Class and superclass level classification of unidentified compounds from UHPLC-MS/MS analysis of control and glyphosate-treated GR-and GS-biotype of *Amaranthus palmeri*.

**Table S5.** Differential Expression analysis of Transcriptome of glyphosate treated GS *Amaranthus palmeri*. Pairwise comparisons were performed using edgeR and mutliple comparisons were controlled using false discovery rate (FDR) approach. Transcripts with p<0.05, FDR <0.05 and log fold change threshold > |1| were considered significant.

**Table S6.** Differential Expression analysis of Transcriptome of glyphosate treated GR *Amaranthus palmeri*. Pairwise comparisons were performed using edgeR and mutliple comparisons were controlled using false discovery rate (FDR) approach. Transcripts with p<0.05, FDR <0.05 and log fold change threshold > |1| were considered significant.

**Table S7.** Pairwise comparison results of proteome of glyphosate-treated GS and GR *Amaranthus palmeri* compared to respective controls. Pairwise comparisons were performed using parametric t-test and mutliple comparisons were controlled using two-stage false discovery rate (FDR) approach of Benjamini, Krieger, and Yekutieli. Proteins with p<0.05, FDR <0.05 and log fold change threshold > |1| were considered significant.

**Table S8.** PTR of glyphosate-and control-treated GS and GR *Amaranthus palmeri*. The statistical analysis for testing significant differences in PTR of different treatments was performed by pairwise comparisons (parametric t-test) and mutliple comparisons were controlled using two-stage false discovery rate (FDR) approach ofBenjamini, Krieger, and Yekutieli.

**Table S9.** Gene Ontology (GO) enrichment analysis for differentially expressed transcripts and proteins in glyphosate-treated GR - and GS-*Amaranthus palmeri*. GO terms with p-value <0.05 and FDR < 0.05 were considered significantly enriched.

**Table S10.** Pairwise comparison results of identified metabolites from GC-QToF analysis of glyphosate-treated GS and GR *Amaranthus palmeri* compared to respective controls. Pairwise comparisons were performed using parametric t-test and mutliple comparisons were controlled using two-stage false discovery rate (FDR) approach of Benjamini, Krieger, and Yekutieli. Metabolites with p<0.05, FDR <0.05 and log fold change threshold > |1| were considered significant.

**Table S11.** Pairwise comparison results of metabolites from UHPLC-MS/MS analysis of glyphosate-treated GS and GR *Amaranthus palmeri* compared to respective controls. Pairwise comparisons were performed using parametric t-test and mutliple comparisons were controlled using two-stage false discovery rate (FDR) approach of Benjamini, Krieger, and Yekutieli. Metabolites with p<0.05, FDR <0.05 and log fold change threshold > |1| were considered significant.

**Table S12.** Paintomics pathways analysis of glyphosate-treated GS (*Amaranthus palmeri*) compared to respective controls.

**Table S13.** Paintomics pathways analysis of glyphosate-treated GR (*Amaranthus palmeri*) compared to respective controls.

## REFERENCES

Bai Y, Kissoudis C, Yan Z, Visser RGF, van der Linden G. 2018. Plant behaviour under combined stress: tomato responses to combined salinity and pathogen stress. The Plant Journal 93: 781–793.

Bathke J, Konzer A, Remes B, McIntosh M, Klug G. 2019. Comparative analyses of the variation of the transcriptome and proteome of Rhodobacter sphaeroides throughout growth. BMC Genomics 20: 1–13.

Benjamini Y, Krieger AM, Yekutieli D. 2006. Adaptive linear step-up procedures that control the false discovery rate. Biometrika 93: 491–507.

Böcker S, Letzel MC, Lipták Z, Pervukhin A. 2009. SIRIUS: decomposing isotope patterns for metabolite identification. Bioinformatics 25: 218–224.

Böcker S, Dührkop K. 2016. Fragmentation trees reloaded. Journal of Cheminformatics 8: 5.

Cañal MJ, Sánchez Tamés R, Fernádez B. 1990. Glyphosate action on abscisic acid levels, stomatal response and electrolyte leakage in yellow nutsedge leaves. Plant Physiology and Biochemistry 28: 215–220.

Clements DR, Ditommaso A. 2011. Climate change and weed adaptation: can evolution of invasive plants lead to greater range expansion than forecasted? Weed Research 51: 227–240.

Culpepper AS, Grey TL, Vencill WK, Kichler JM, Webster TM, Brown SM, York AC, Davis JW, Hanna WW. 2006. Glyphosate-resistant Palmer amaranth (*Amaranthus palmeri*) confirmed in Georgia. Weed Science 54: 620–626.

de Godoy LMF, Olsen J v., Cox J, Nielsen ML, Hubner NC, Fröhlich F, Walther TC, Mann M. 2008. Comprehensive mass-spectrometry-based proteome quantification of haploid versus diploid yeast. Nature 455: 1251–1254.

de Visser R, Spitters CJT, Bouma TJ. 1992 Energy cost of protein turnover: theoretical calculation and experimental estimation from regression of respiration on protein concentration of full-grown leaves. *Molecular*, Biochemical and Physiological Aspects of Plant Respiration, 493-508.

Délye C, Jasieniuk M, le Corre V. 2013. Deciphering the evolution of herbicide resistance in weeds. Trends in Genetics 29: 649–658.

Djoumbou Feunang Y, Eisner R, Knox C, et al. 2016 ClassyFire: Automated chemical classification with a comprehensive, computable taxonomy. Journal of Cheminformatics 8: 61.

Dührkop K, Shen H, Meusel M, Rousu J, Böcker S. 2015. Searching molecular structure databases with tandem mass spectra using CSI:FingerID. Proceedings of the National Academy of Sciences of the United States of America 112: 12580–5.

Dührkop K, Fleischauer M, Ludwig M, Aksenov AA, Melnik A v., Meusel M, Dorrestein PC, Rousu J, Böcker S. 2019. SIRIUS 4: a rapid tool for turning tandem mass spectra into metabolite structure information. Nature Methods 16: 299–302.

Dührkop K, Nothias L-F, Fleischauer M, Reher R, Ludwig M, Hoffmann MA, Petras D, Gerwick WH, Rousu J, Dorrestein PC, et al. 2021. Systematic classification of unknown metabolites using high-resolution fragmentation mass spectra. Nature Biotechnology 39: 462– 471.

Duke SO, Powles SB. 2008. Glyphosate: a once-in-a-century herbicide. Pest Management Science 64: 319–325.

Dyer WE. 2018. Stress-induced evolution of herbicide resistance and related pleiotropic effects. Pest Management Science 74: 1759–1768.

Farooq U, Rehman A, Ashraf MA, Rasheed R, Shahid M, Ali S, Sarker PK. 2025. Taurine priming improves redox balance, osmotic adjustment, and nutrient acquisition to lessen phytotoxic effects of neutral and alkaline salts on pea (Pisum sativum L.). Plant Signal Behav. 20(1): 2480224.

Faus I, Zabalza A, Santiago J, Nebauer SG, Royuela M, Serrano R, Gadea J. 2015. Protein kinase GCN2 mediates responses to glyphosate in Arabidopsis. BMC Plant Biology 15: 1–12.

Fernández-Escalada M, Gil-Monreal M, Zabalza A, Royuela M. 2016. Characterization of the *Amaranthus palmeri* physiological response to glyphosate in susceptible and resistant populations. Journal of Agricultural and Food Chemistry 64: 95–106.

Fernández-Escalada M, Zulet-González A, Gil-Monreal M, Zabalza A, Ravet K, Gaines T, Royuela M. 2017. Effects of EPSPS Copy Number Variation (CNV) and glyphosate application on the aromatic and branched chain amino acid synthesis pathways in *Amaranthus palmeri*. Frontiers in Plant Science 8: 1970.

Fernández-Escalada M, Zulet-González A, Gil-Monreal M, Royuela M, Zabalza A. 2019. Physiological performance of glyphosate and imazamox mixtures on *Amaranthus palmeri* sensitive and resistant to glyphosate. Scientific Reports 9: 18225.

Fournier ML, Paulson A, Pavelka N, Mosley AL, Gaudenz K, Bradford WD, Glynn E, Li H, Sardiu ME, Fleharty B, et al. 2010. Delayed correlation of mRNA and protein expression in rapamycin-treated cells and a role for ggc1 in cellular sensitivity to rapamycin. Molecular & Cellular Proteomics 9: 271.

Fuchs B, Saikkonen K, Helander M. 2021. Glyphosate-modulated biosynthesis driving plant defense and species interactions. Trends in Plant Science 26: 312–323.

Gaines TA, Zhang W, Wang D, Bukun B, Chisholm ST, Shaner DL, Nissen SJ, Patzoldt WL, Tranel PJ, Culpepper AS, et al. 2010. Gene amplification confers glyphosate resistance in *Amaranthus palmeri*. Proceedings of the National Academy of Sciences of the United States of America 107: 1029–1034.

Gaines TA, Shaner DL, Ward SM, Leach JE, Preston C, Westra P. 2011. Mechanism of resistance of evolved glyphosate-resistant Palmer amaranth (*Amaranthus palmeri*). Journal of Agricultural and Food Chemistry 59: 5886–5889.

Gaines TA, Duke SO, Morran S, Rigon CAG, Tranel PJ, Küpper A, Dayan FE. 2020. Mechanisms of evolved herbicide resistance. Journal of Biological Chemistry 295: 10307– 10330.

Gimenez E, Salinas M, Manzano-Agugliaro F. 2018. Worldwide research on plant defense against biotic stresses as improvement for sustainable agriculture. Sustainability 10: 391.

Gomes MP, Smedbol E, Chalifour A, Hénault-Ethier L, Labrecque M, Lepage L, Lucotte M, Juneau P. 2014. Alteration of plant physiology by glyphosate and its by-product aminomethylphosphonic acid: an overview. Journal of Experimental Botany 65: 4691–4703.

Gomes MP, le Manac’h SG, Maccario S, Labrecque M, Lucotte M, Juneau P. 2016a. Differential effects of glyphosate and aminomethylphosphonic acid (AMPA) on photosynthesis and chlorophyll metabolism in willow plants. Pesticide Biochemistry and Physiology 130: 65–70.

Gomes MP, Juneau P. 2016b. Oxidative stress in Duckweed (*Lemna minor* L.) induced by glyphosate: Is the mitochondrial electron transport chain a target of this herbicide? Environmental Pollution 218: 402–409.

Gomes MP, le Manac’h SG, Hénault-Ethier L, Labrecque M, Lucotte M, Juneau P. 2017. Glyphosate-dependent inhibition of photosynthesis in willow. Frontiers in Plant Science 8: 207.

Haak DC, Fukao T, Grene R, Hua Z, Ivanov R, Perrella G, Li S. 2017. Multi-level Regulation of Abiotic Stress Responses in Plants. Frontiers in Plant Science 8: 1564.

Heap I, Duke SO. 2018. Overview of glyphosate-resistant weeds worldwide. Pest Management Science 74: 1040–1049.

Heap, I. The International Herbicide-Resistant Weed Database. Online. Saturday, July 12, 2025. Available www.weedscience.org

Hoffmann MA, Nothias L-F, Ludwig M, Fleischauer M, Gentry EC, Witting M, Dorrestein PC, Dührkop K, Böcker S. 2021. High-confidence structural annotation of metabolites absent from spectral libraries. Nature Biotechnology, DOI: 10.1038/s41587-021-01045-9.

Ishihara H, Moraes TA, Pyl ET, Schulze WX, Obata T, Scheffel A, Fernie AR, Sulpice R, Stitt M. 2017. Growth rate correlates negatively with protein turnover in Arabidopsis accessions. The Plant Journal 91: 416–429.

Jamil IN, Remali J, Azizan KA, Nor Muhammad NA, Arita M, Goh H-H, Aizat WM. 2020. Systematic multi-omics integration (MOI) approach in plant systems biology. Frontiers in Plant Science 11: 944.

Jiang LX, Jin LG, Guo Y, Tao B, Qiu LJ. 2013. Glyphosate effects on the gene expression of the apical bud in soybean (*Glycine max*). Biochemical and Biophysical Research Communications 437: 544–549.

Kim G, Clarke CR, Larose H, Tran HT, Haak DC, Zhang L, Askew S, Barney J, Westwood JH. 2017. Herbicide injury induces DNA methylome alterations in Arabidopsis. PeerJ 5: e3560.

Koo D, Molin WT, Saski CA, Jiang J, Putta K, Jugulam M, Friebe B, Gill BS. 2018. Extrachromosomal circular DNA-based amplification and transmission of herbicide resistance in crop weed *Amaranthus palmeri*. Proc. Natl. Acad. Sci. U.S.A. 115(13): 3332–3337.

Kumar M, Brar A, Yadav M, Chawade A, Vivekanand V, Pareek N. 2018 Chitinases-potential candidates for enhanced plant resistance towards fungal pathogens. Agriculture 8: 88.

Liu T, Salguero P, Petek M, Martinez-Mira C, Balzano-Nogueira L, Ramšak Ž, McIntyre L, Gruden K, Tarazona S, Conesa A. 2022. PaintOmics 4: new tools for the integrative analysis of multi-omics datasets supported by multiple pathway databases. Nucleic Acids Research 50: 551– 559.

Ludwig M, Nothias L-F, Dührkop K, Koester I, Fleischauer M, Hoffmann MA, Petras D, Vargas F, Morsy M, Aluwihare L, et al. 2020. Database-independent molecular formula annotation using Gibbs sampling through ZODIAC. Nature Machine Intelligence 2: 629–641.

Maeda H, Dudareva N. 2012. The shikimate pathway and aromatic amino acid biosynthesis in plants. Annual Review of Plant Biology 63: 73–105.

Maroli AS, Nandula VK, Dayan FE, Duke SO, Gerard P, Tharayil N. 2015. Metabolic profiling and enzyme analyses indicate a potential role of antioxidant systems in complementing glyphosate resistance in an *Amaranthus palmeri* biotype. Journal of Agricultural and Food Chemistry 63: 9199–9209.

Maroli A, Nandula V, Duke S, Tharayil N. 2016. Stable isotope resolved metabolomics reveals the role of anabolic and catabolic processes in glyphosate-induced amino acid accumulation in *Amaranthus palmeri* biotypes. Journal of Agricultural and Food Chemistry 64: 7040–7048.

Matsuo T, Tsugawa H, Miyagawa H, Fukusaki E. 2017. Integrated strategy for unknown EI–MS identification using quality control calibration curve, multivariate analysis, EI–MS spectral database, and retention index prediction. Analytical Chemistry 89: 6766–6773.

Meena KK, Sorty AM, Bitla UM, Choudhary K, Gupta P, Pareek A, Singh DP, Prabha R, Sahu PK, Gupta VK, et al. 2017. Abiotic stress responses and microbe-mediated mitigation in plants: the omics strategies. Frontiers in Plant Science 0: 172.

Mergner J, Frejno M, List M, Papacek M, Chen X, Chaudhary A, Samaras P, Richter S, Shikata H, Messerer M, et al. 2020. Mass-spectrometry-based draft of the Arabidopsis proteome. Nature 579: 409–414.

Misra BB, Langefeld C, Olivier M, Cox LA. 2019. Integrated omics: Tools, advances and future approaches. Journal of Molecular Endocrinology 62: 21–45.

Nandula VK, Reddy KN, Koger CH, Poston DH, Rimando AM, Duke SO, Bond JA, Ribeiro DN. 2012. Multiple resistance to glyphosate and pyrithiobac in Palmer amaranth (*Amaranthus palmeri*) from Mississippi and response to flumiclorac. Weed Science 60: 179–188.

Nishizawa A, Yabuta Y, Shigeoka S. 2008. Galactinol and Raffinose Constitute a Novel Function to Protect Plants from Oxidative Damage. Plant Physiology 147(3), 1251–1263.

Oh DH, Dassanayake M, Bohnert HJ, Cheeseman JM. 2012. Life at the extreme: lessons from the genome. Genome Biology 13: 1–9.

Orcaray L, Zulet A, Zabalza A, Royuela M. 2012. Impairment of carbon metabolism induced by the herbicide glyphosate. Journal of Plant Physiology 169: 27–33.

Patti GJ, Yanes O, Siuzdak G. 2012. Metabolomics: the apogee of the omics trilogy. Nature Reviews Molecular Cell Biology 13: 263–269.

Sandhu PK, Leonard E, Nandula V, Tharayil N. 2023. Global Metabolome of Palmer Amaranth (*Amaranthus palmeri*) Populations Highlights the Specificity and Inducibility of Phytochemical Responses to Abiotic Stress. Journal of Agricultural and Food Chemistry 71: 3518–3530.

Schwartz TS. 2020. The promises and the challenges of integrating multi-omics and systems biology in comparative stress biology. Integrative and Comparative Biology 60: 89–97.

Schymanski EL, Jeon J, Gulde R, Fenner K, Ruff M, Singer HP, Hollender, J. 2014. Identifying small molecules via high resolution mass spectrometry: Communicating confidence. Environmental Science and Technology 48: 2097−2098.

Soda N, Wallace S, Karan R. 2015. Omics study for abiotic stress responses in plants. Advances in Plants & Agriculture Research 2.

Srivastava V, Obudulu O, Bygdell J, Löfstedt T, Rydén P, Nilsson R, Ahnlund M, Johansson A, Jonsson P, Freyhult E, et al. 2013. OnPLS integration of transcriptomic, proteomic and metabolomic data shows multi-level oxidative stress responses in the cambium of transgenic hipI-superoxide dismutase Populus plants. BMC Genomics 14: 1–16.

Sun S, Zhou J. 2018. Molecular mechanisms underlying stress response and adaptation. Thoracic Cancer 9: 218–227.

Tsugawa H, Tsujimoto Y, Arita M, Bamba T, Fukusaki E. 2011a. GC/MS based metabolomics: development of a data mining system for metabolite identification by using soft independent modeling of class analogy (SIMCA). BMC bioinformatics 12: 131.

Tsugawa H, Bamba T, Shinohara M, Nishiumi S, Yoshida M, Fukusaki E. 2011b. Practical non-targeted gas chromatography/mass spectrometry-based metabolomics platform for metabolic phenotype analysis. Journal of bioscience and bioengineering 112: 292–8.

Vengavasi K, Kumar A, Pandey R. 2016. Transcript abundance, enzyme activity and metabolite concentration regulates differential carboxylate efflux in soybean under low phosphorus stress. Indian Journal of Plant Physiology 21: 179–188.

Vila-Aiub MM, Neve P, Powles SB. 2009. Fitness costs associated with evolved herbicide resistance alleles in plants. New Phytologist 184: 751–767.

Vila-Aiub MM, Yu Q, Powles SB. 2019. Do plants pay a fitness cost to be resistant to glyphosate? New Phytologist 223: 532–547.

Wang P, Hsu CC, Du Y, Zhu P, Zhao C, Fu X, Zhang C, Paez JS, Macho AP, Andy Tao W, et al. 2020. Mapping proteome-wide targets of protein kinases in plant stress responses. Proceedings of the National Academy of Sciences of the United States of America 117: 3270– 3280.

Ward SM, Webster TM, Steckel LE. 2013. Palmer amaranth (*Amaranthus palmeri*): A review. Weed Technology 27: 12–27.

Young MD, Wakefield MJ, Smyth GK, Oshlack A 2010. Gene ontology analysis for RNA-seq: accounting for selection bias. Genome Biology 11: R14.

Zhang L, Du L, Poovaiah BW. 2014. Calcium signaling and biotic defense responses in plants. Plant Signaling & Behavior 9: e973818.

