## Supplementary Figures for "Integrated multi-omics analysis reveals divergent molecular responses in Palmer amaranth (*Amaranthus palmeri*) biotypes susceptible and resistant to glyphosate"

**Figure S1. PLS-DA plot of first two components from PLS-DA of transcriptomics, proteomics, primary (GC-QToF), and secondary metabolomics (UHPLC-MS/MS) data.** Different treatments are represented by different colors and ellipses represent 95% confidence interval.

**Figure S2. PTR Density plot of the protein-transcript pairs.** Density plot of protein-mRNA ratio (PTR) for the protein with corresponding transcript abundance in control and glyphosate-treated GS and GR biotype of *Amaranthus palmer*i. The straight vertical line represents the median values for each treatment.

**Figure S3. Identified terpenoids of the present study.** Asterisk denotes significant difference ($p<0.05$, FDR<0.05, log2 fold change >|1|).

**Figure S4: Effect of glyphosate on transcripts and proteins of aromatic amino acid biosynthesis pathway in GS and GR *Amaranthus palmeri*.** Heatmap displaying the $\log_2$ fold change in the abundance of transcripts and proteins of aromatic amino acid biosynthesis pathway. Column C is the $\log_2$ fold change of GR control vs GS control. GS and GR respectively denote the $\log_2$ fold change of glyphosate treatment with respective control for GS and GR-biotypes. First three columns are for transcript abundance and the next three for protein abundance. Asterisk denotes significant differences at $p<0.05$, FDR <0.05, and $\log_2$ fold change >|1|.

**Figure S5. Agarose gel electrophoresis of isolated total RNA from leaves of *Amaranthus palmeri*.** RNA bands (28S, 18S, 5S) are indicated by arrows. Samples with a ratio of 2:1 for 28S:18S bands are considered good quality. The ID on top of each gel column represents sample ID. For details about samples, refer to table S1.

**Figure S6. Total soluble protein in leaves of *Amaranthus palmeri*.**

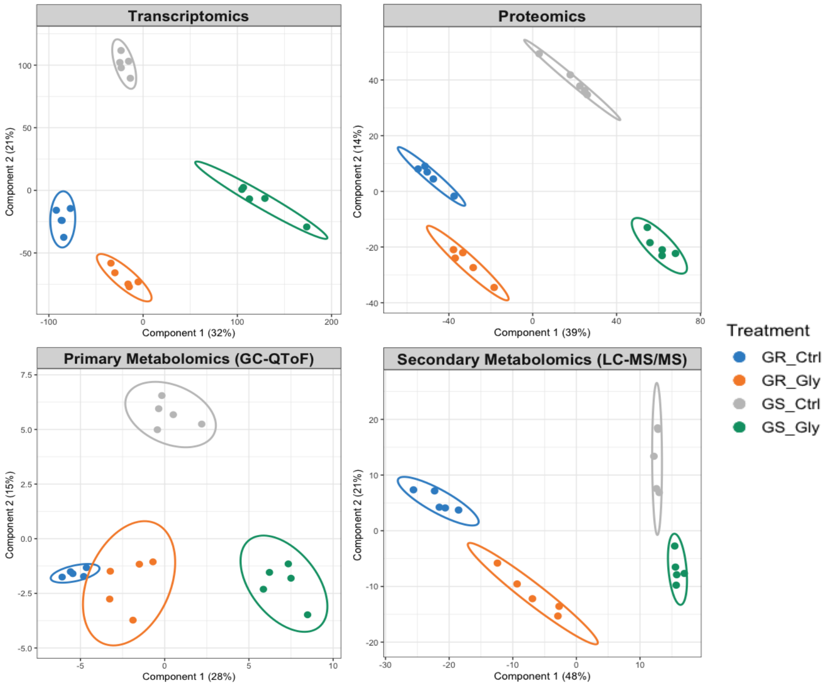


**Figure S1. PLS-DA plot of first two components from PLS-DA of transcriptomics, proteomics, primary (GC-QToF), and secondary metabolomics (UHPLC-MS/MS) data.** Different treatments are represented by different colors, and ellipses represent 95% confidence intervals.

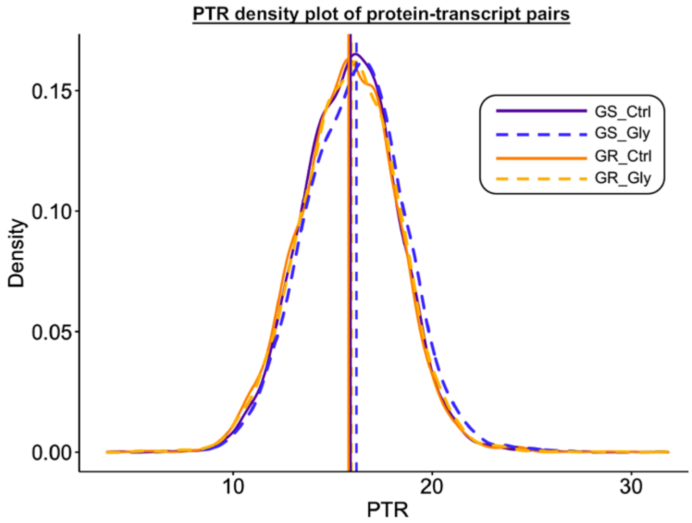


**Figure S2. PTR Density plot of the protein-transcript pairs.** Density plot of protein-mRNA ratio (PTR) for the protein with corresponding transcript abundance in control and glyphosate-treated GS and GR biotype of *Amaranthus palmeri*. The straight vertical line represents the median values for each treatment.

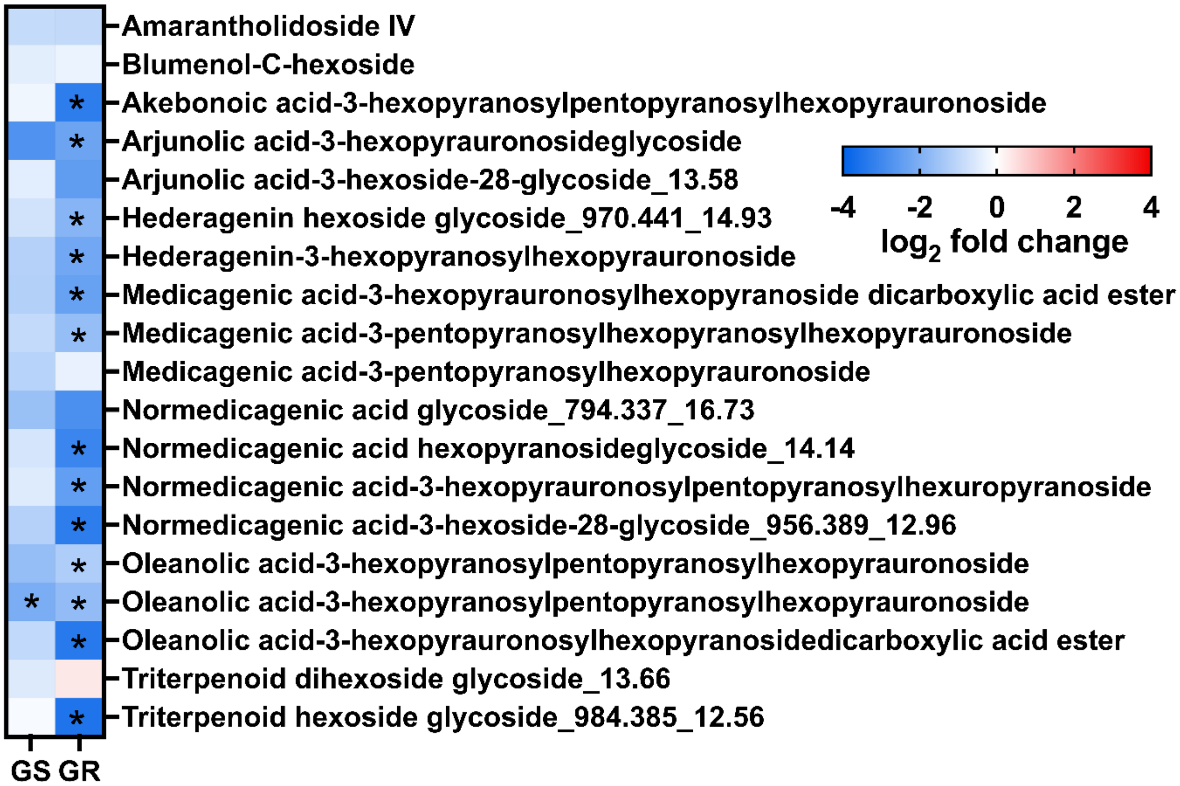


**Figure S3. Identified terpenoids of the present study.** Asterisk denotes significant difference ($p<0.05$, FDR<0.05, log2 fold change >|1|).

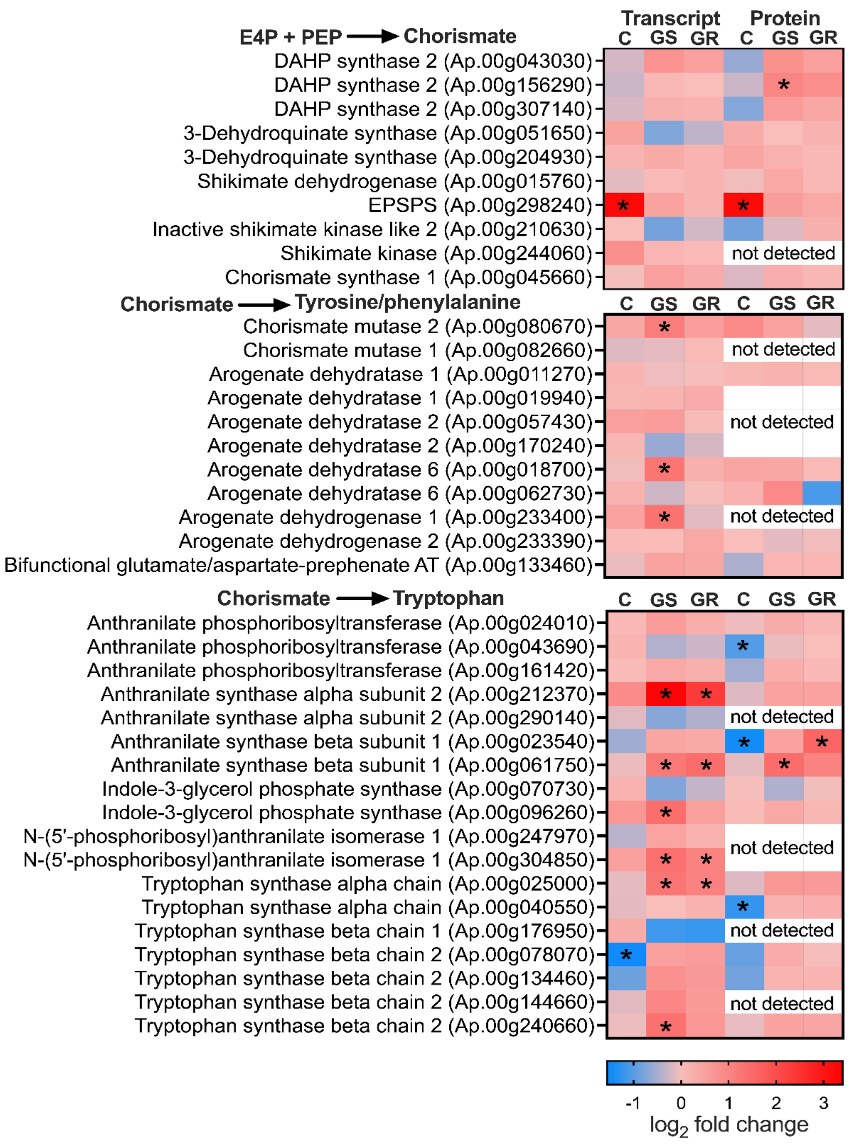


**Figure S4. Effect of glyphosate on transcripts and proteins of aromatic amino acid biosynthesis pathway in GS and GR *Amaranthus palmeri*.** Heatmap displaying the $\log_2$ fold change in the abundance of transcripts and proteins of aromatic amino acid biosynthesis pathway. Column C is the $\log_2$ fold change of GR control vs GS control. GS and GR respectively denote the $\log_2$ fold change of glyphosate treatment with respective control for GS and GR-biotypes. First three columns are for transcript abundance and the next three for protein abundance. Asterisk denotes significant differences at $p<0.05$, FDR $<0.05$, and $\log_2$ fold change $>|1|$.

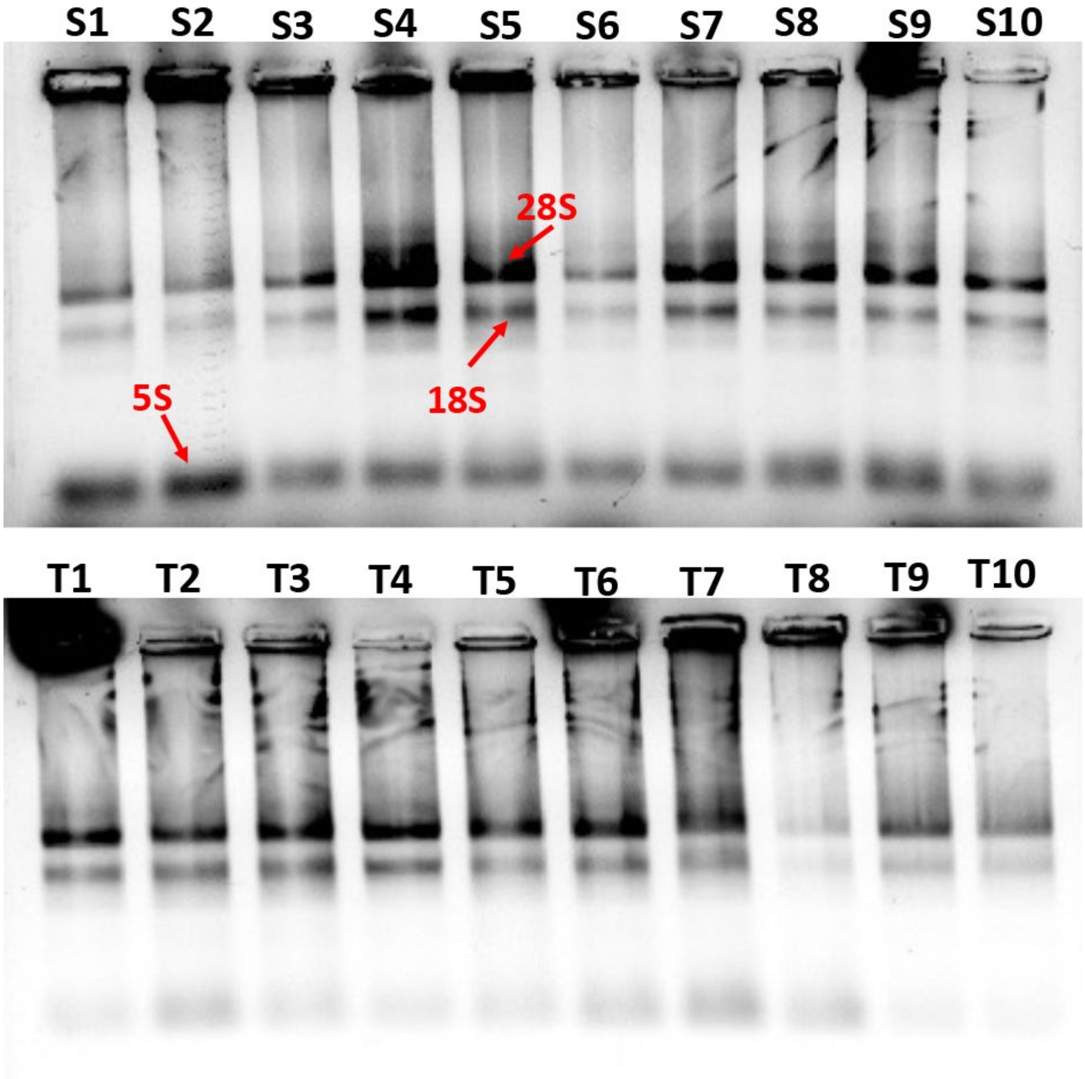


**Figure S5. Agarose gel electrophoresis of isolated total RNA from leaves of *Amaranthus palmeri*.** RNA bands (28S, 18S, 5S) are indicated by arrows. Samples with a ratio of 2:1 for 28S:18S bands are considered good quality. The ID on top of each gel column represents sample ID. For details about samples, refer to table S1.

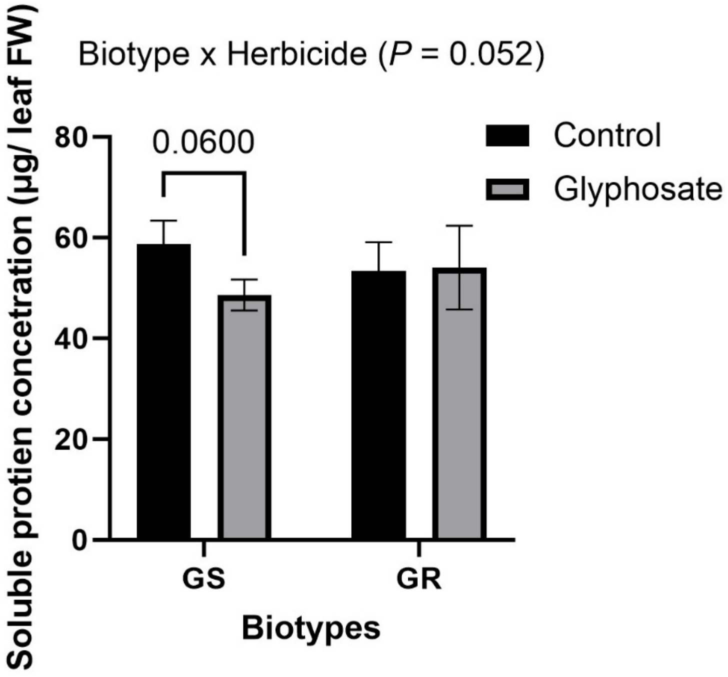


**Figure S6. Total soluble protein in leaves of *Amaranthus palmeri*.**
